# Kinesin-1 conformational dynamics are controlled by a cargo-sensitive TPR switch

**DOI:** 10.1101/2025.04.08.647705

**Authors:** Shivam Shukla, Jessica A. Cross, Monika Kish, Sathish K.N. Yadav, Johannes F. Weijman, Laura O’Regan, Judith Mantell, Ufuk Borucu, Xiyue Leng, Christiane Schaffitzel, Jonathan J. Phillips, Derek N. Woolfson, Mark P. Dodding

## Abstract

Kinesin-1 is a dynamic heterotetrameric assembly of two heavy and two light chains (KHC and KLC) that mediates microtubule-based intracellular transport of many different cargoes. The complex adopts a compact, autoinhibited state that is activated by cargo-adaptor proteins containing specific short linear peptide motifs (SLiMs). These motifs interact with the tetratricopeptide repeat (TPR) domains of the KLCs. The mechanism coupling SLiM recognition to activation-associated conformational changes in the complex is unknown. Here we combine protein design, computational modelling, biophysical analysis, and electron microscopy to examine the structural and mechanistic consequences of SLiM binding to the KLC-TPR domain within the complete heterotetrameric holoenzyme. We show that coiled coil 1 (CC1) of the KHC docks KLC TPR domains in the autoinhibited complex, forming the ‘shoulder’ feature observed in electron microscopy. Disrupting this interaction or binding an activating SLiM dislocates the TPR shoulder, freeing the motor domains and promoting transition between its closed, inactive, and open states. Opening the kinesin-1 complex facilitates binding to the microtubule-associated kinesin-1 cofactor, MAP7. Therefore, cargo-mediated dislocation of the TPR shoulder serves as a key initial step in kinesin-1 activation, allosterically linking cargo binding to motor dynamics.

## Introduction

Intracellular transport relies on the conformational dynamics of cytoskeletal motor proteins. One such process involves ATPase-driven binding and movement along cytoskeletal tracks. Another, less-understood process, involves shape changes in motor complexes, which regulate movement in response to inputs like cargo or regulatory protein binding, ensuring precise transport control (*1–4*). Conformational transitions between a compact inhibited state and an open active state are essential for the regulation of kinesin-1, a ubiquitous and prototypic family of microtubule motors involved in transporting proteins, ribonucleoproteins, vesicles, and organelles in cells (*1, 2, 5–8*). Kinesin-1 is also hijacked by pathogens during infection, and its dysregulation is linked to neurological disorders (*9, 10*).

Heterotetrameric kinesin-1 consists of two kinesin heavy chains (KHCs) and two kinesin light chains (KLCs). In mammals, KHCs are encoded by three paralogs—KIF5A, KIF5B, and KIF5C—while KLCs are encoded by four paralogs (KLC1–KLC4) (*11–14*). The KHCs have ATPase motor domains at the amino terminus, followed by coiled-coil domains (CC0–CC4) that mediate KHC dimerization, CC0 forms the neck between the amino-terminal motor domains and the coiled-coil stalk which comprises CC1 – CC4. The carboxy-terminus contains an unstructured domain with the autoinhibitory ’IAK’ motif, which regulates ATPase activity in the compact state, but is not essential for its formation (*15–23*). The KLCs, which also help to regulate kinesin-1 activity, have a coiled-coil domain that binds to CC3 of the KHCs, followed by an unstructured linker to a tetratricopeptide repeat (TPR) domain critical for cargo binding (*2, 24–26*) (**Fig. 1A)**. In several KLC isoforms, the TPR domain is followed by an intrinsically disordered domain, which includes a membrane-binding amphipathic helix that can stabilise interactions with membranous cargoes (*27–29*).

**Figure 1.**
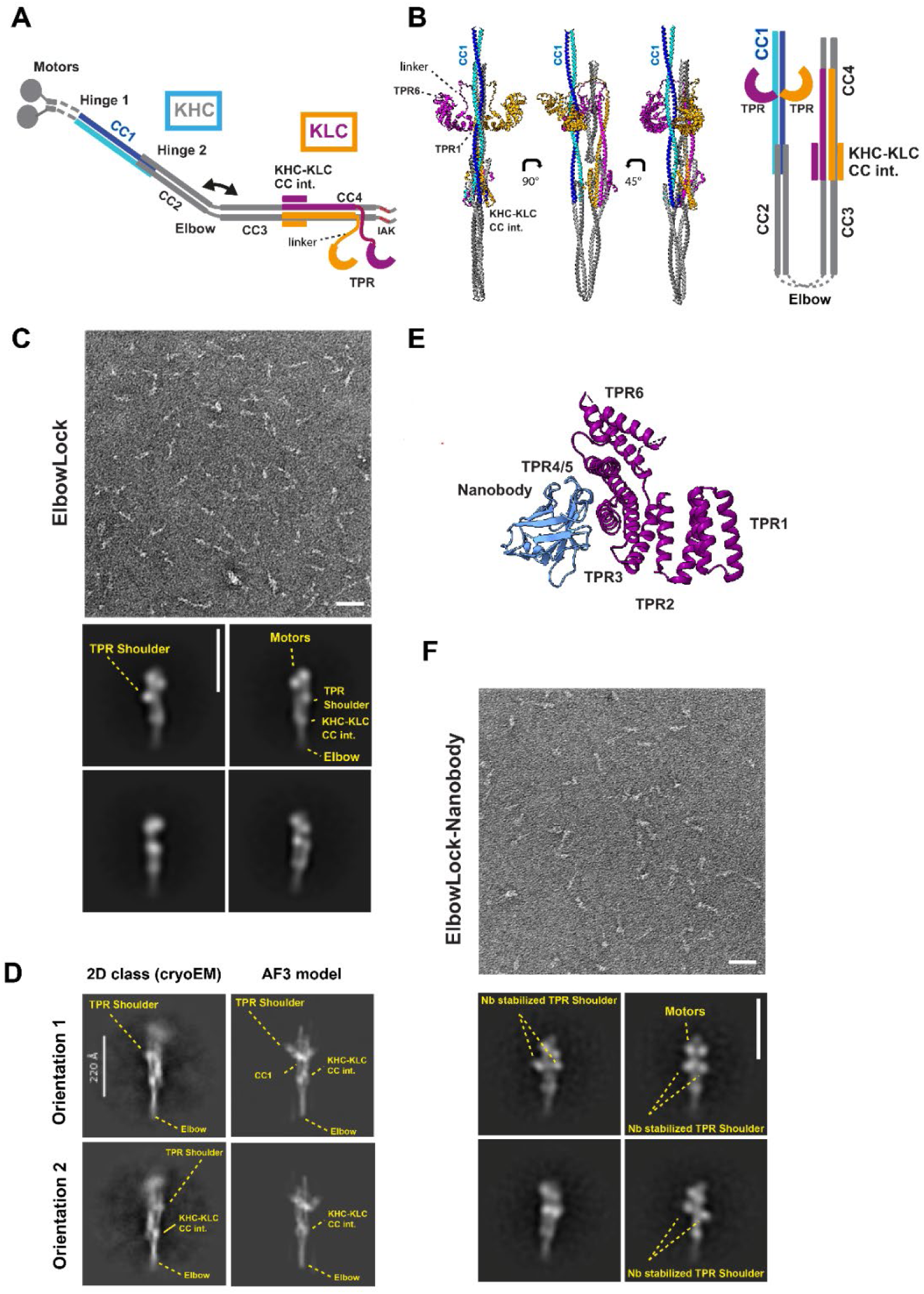
Computational modelling and single particle EM analysis identify a TPR docking site (TDS) on CC1. (A) Schematic representation of heterotetramer kinesin-1 in an open conformation. (B) AF3 model in 3 orientations alongside a schematic showing KHC(KIF5C) coiled-coil assembly with KLCs. KHCs are shown in blue and cyan, KLCs in orange and purple. The complete AF3 output, including models for the full tetramer, and sequences used are described in Figure S1. Schematic shows predicted domain positioning, omitting linker sequences and separating the KHC coiled-coils for clarity (C) Representative NS electron micrograph (from 3 independent experiments) showing ElbowLock complexes (scale bar is 50 nm) with selected 2D classes from NS-EM (scale bar is 26 nm). (D) Reference-free 2D class averages showing two orientations of complete heterotetrameric kinesin-1 complexes, compared to low-pass filtered back-projection of the AF3 model (scale bar is 22 nm). (E) Structure of the nanobody-TPR complex highlighting binding on exterior of TPRs 4 and 5 (pdb:6fuz). Cargo JIP1 Y-acidic peptide was removed for clarity.(F) Representative NS electron micrograph showing ElbowLock-nanobody complexes (scale bar is 50 nm) with selected 2D classes from NS-EM (scale bar is 26 nm).

Kinesin-1 activity is regulated by its conformational state (*6–8, 24, 30, 31*). Recent studies using negative stain electron microscopy (NS-EM), AlphaFold2 modelling, small-angle X-ray scattering (SAXS) and cross-linking mass spectrometry have revealed the architecture of the compact, autoinhibited state, named the “lambda particle” (*22, 32, 33*). Key features of this particle include the ATPase motor domains, forming the “head,” and a flexion point in the coiled-coil scaffold between CC2 and CC3—the “elbow”—that enables the autoinhibited conformation to form (*22, 32, 33*). This flexion is functionally significant, as the complex can be activated or stabilized in its inhibited state through targeted mutations or binding of *de novo* designed peptide-based activators (*32, 34*). NS-EM and SAXS also reveal a “shoulder” near the motor domains. Domain-deletion experiments indicate that the shoulder is formed by at least one KLC TPR domain (*32*). Cross-linking mass spectrometry, NS-EM, and AlphaFold2 modelling suggest that the motor domains invert relative to CC1, binding back upon it and interact with CC4 in a hierarchical inhibition mechanism (*22, 33*) and propose many possible contacts between the KLC-TPR domains with the motor domains and coiled-coil body (*22*). Key features of the inhibited complex appear to be conserved across the closely related mammalian KHC paralogues (*22, 32, 33*).

Understanding the position and function of TPR domains in the inhibited state is crucial for uncovering how kinesin-1 is activated, as these domains provide binding sites for activating cargo adaptors with W-acidic SLiMs (*2, 35, 36*). W-acidic motifs bind the concave groove of the TPR superhelix, increase its curvature, and displace an autoregulatory interaction with the CC-TPR linker region to modulate KLC conformational state (*24*). This molecular switch is also the target for the kinesin-1 activating small-molecule kinesore (*37*). Adaptors carrying W-acidic motifs include the lysosomal adaptor SKIP (*35, 38, 39*), the nuclear envelope adaptors Nesprin-2/4 (*40, 41*), and the amyloid-precursor protein (APP) vesicle adaptor calsyntenin-1 (*42–44*), and others (*2*). Recently, we have designed a small-peptide ligand, KinTag, for the TPR binding site. This combines features of natural micromolar affinity W-acidic and related Y-acidic motifs, to give high-affinity (low nanomolar) binding through simultaneous occupation of the W-, Y- and F- pockets on the TPR receptor (*26, 45, 46*). Notably, KinTag drives changes in TPR conformation like the natural ligands. Therefore, KinTag provides a useful tool to probe the mechanistic consequences of stable motor-cargo interactions, which in the natural system, are also enabled by other co-operative protein-protein and protein-lipid interactions (*27, 47*). Importantly, when fused to an integral membrane protein, KinTag, or its parental W-acidic sequence, is sufficient to drive kinesin-organelle recruitment and axonal transport (*36, 42, 48, 49*). Thus, structural and biophysical analysis of isolated TPR domains in vitro and functional analysis of organelle transport in cells demonstrates key role for SLiM-TPR binding in motor activation and has defined some important components of the pathway. However, the mechanism that couples SLiM cargo recognition to activation-linked conformational change in the complete kinesin-1 holoenzyme remains unknown.

To define this mechanism, we use protein design and engineering approaches to overcome intrinsic conformational dynamics and low-affinity motor-cargo interactions to stabilize intermediates along the kinesin-1 inhibition-activation pathway, enabling their biochemical characterization and visualization. We show that the KLC TPR domains dock onto CC1 in the autoinhibited complex to form the shoulder. Disrupting this interaction or binding an activating SLiM dislocates the shoulder. This makes the motor domains more accessible and dynamic as judged by hydrogen-deuterium exchange measurements in the open, active, and cargo-bound states compared with the closed, inactive form. Opening the complex also promotes interaction with the microtubule-associated cofactor, MAP7. Therefore, cargo-mediated dislocation of the TPR shoulder serves as a key step in kinesin-1 activation, allosterically linking cargo recognition to conformational dynamics.

## Results

### Coiled coil 1 (CC1) provides a TPR docking site (TDS) in the autoinhibited kinesin-1 complex

To begin, we sought to clarify the position of the TPR domains in the lambda particle by modelling their potential for associations using AlphaFold3 (AF3). AF3-generated assemblies of the heterotetrameric complexes of the KHC coiled coils (KIF5C) and KLC (KLC1A), that included the TPR domains (KIF5C/KLC1A) yielded models that were invariably folded-over at the elbow, with a global coiled-coil architecture consistent with our previous work (*32, 34*) **(Fig. 1B, Fig. S1A)**. Interestingly, the two TPR domains were confidently predicted to bind with C2 symmetry on CC1 ‘end on’ via the first repeats (TPR1) at a position concurrent with the shoulder observed in NS-EM. This TPR conformation closely resembles their known mode of binding to the coiled-coil leucine zipper motif of the cargo adaptor JIP3 (*50*). The concave cargo binding surface on the TPR domain that recognises SLiM adaptors was typically occupied by the linker region that connects the TPR to the KHC-KLC coiled-coil interface (*24*). Models of the complete heterotetramer, including the motor domains were similar. The motor domains were inverted around Hinge 1 enabling interactions between them and the first part of CC1 and the end of CC4 with the neck coiled coil (CC0) projecting away from the motors, in line with recent reports (*22, 33*) **(Fig. S1B,C,D)**.

To test these predictions experimentally, we examined purified kinesin-1 complexes using negative stain electron microscopy (NS-EM) followed by 2D-classification of single particles **(Fig. S2, S3, S4)**. We focused on the ElbowLock background. This KIF5C construct contains a short, 5 amino acid, deletion that restricts flexibility around the elbow and helps maintain particles in their lambda conformation, providing homogenous samples, and facilitating subsequent analysis (*34*). Proteins were also stabilised using the amine-to-amine crosslinker BS3 that was important for achieving reproducibly high-quality samples for imaging. As observed previously for closed wild-type particles, the TPR ‘shoulder’ feature was readily apparent, and typically appeared as a single globular density proximal to the two motor domains (*22, 32*) **(Fig. 1C)**.

To visualise finer structural details, we turned to single-particle cryoEM analysis of frozen-hydrated samples. We were unable to obtain optimal samples suitable for determining the complete structure. Nonetheless, we obtained reference-free 2D class averages that appeared to show full-length ‘side’ views of the complex with clear definition of the elbow, hinge 2, and KHC-KLC (coiled-coil) interface features **(Fig. 1D)**. The motor domains were poorly resolved in these classes, suggesting that the head assembly is somewhat flexible relative to the coiled coil/TPR body. A comparison to low-pass filtered back-projections from the AF3 model (without motor domains) revealed density at a position concurrent with the docked TPR domains **(Fig. 1D)**.

However, TPR domain stoichiometry at the shoulder remained unclear. We reasoned that if the AF3 models were correct then the latter TPR repeats (4–6) would be positioned away from the interface with the CC1. The KLC TPR domains are small and known to be conformationally dynamic, and so this might make it difficult to resolve them in 2D class averages (*24, 35, 45*). To test this hypothesis, we assembled complexes with a nanobody to bind and stabilise TPRs 4 and 5 and add additional mass (13 kDa per TPR) (*36, 45*) **(Fig. 1E, Fig. S2)**. This resulted in NS-EM 2D classes with density extending either side of the coiled-coil scaffold that were fully consistent with the AF3 tetramer models and binding of both TPR domains on CC1 **(Fig. 1E, F)**. There was no evidence of additional density elsewhere in the complex. Thus, AF3 predicts a model for TPR domain association with the coiled-coil scaffold that is supported by electron microscopy experiments. Because we have used a stabilisation approach, we do not exclude the possibility that there may be differences in TPR stability, dynamics and occupancy at the two potential CC1 sites in the absence of stabilisation. i.e. only one TPR may be typically bound at one time, and/or they may have potential for exchange. Hereafter, the predicted <u>T</u>PR <u>d</u>ocking <u>s</u>ite on CC1 is called the TDS.

### Removal of the TDS dislocates the kinesin-1 shoulder

To test the role of the TDS in positioning the KLC TPRs directly, we removed it while preserving the overall length and coiled-coil structure of CC1. To do this, four heptad repeats of CC1 of KIF5C, flanking and incorporating TDS, were replaced with a *de novo*-designed homodimeric coiled coil, CC-Di (*51*). The aim was to retain the dimeric coiled-coil structure and length of CC1 while eliminating side chains that promote its interface with the TPR (ΔTDS) **(Fig. 2A)**. Supporting this, models of ΔTDS complexes using AF3 showed the expected seamless insertion of CCDi into CC1, with displacement of the TPR domains to a variety of different positions, in 5 models, all with high position error with respect to KHC **(Fig. S5)**. We used size-exclusion chromatography (SEC) to further validate our design. ElbowLock proteins elute as a single lambda peak and this elution profile was unchanged by the ΔTDS modification, indicating that complex assembly or folding is not disrupted **(Fig. 2B)**. ElbowLock-ΔTDS complexes were then analyzed by NS-EM and 2D classification (**Fig. 2 C,D)** and quantitatively compared to the parental ElbowLock **(Fig. 2E)**. The shoulder was lost in the ΔTDS background, suggesting that TPR domains had undocked from the core of the complex.

**Figure 2.**
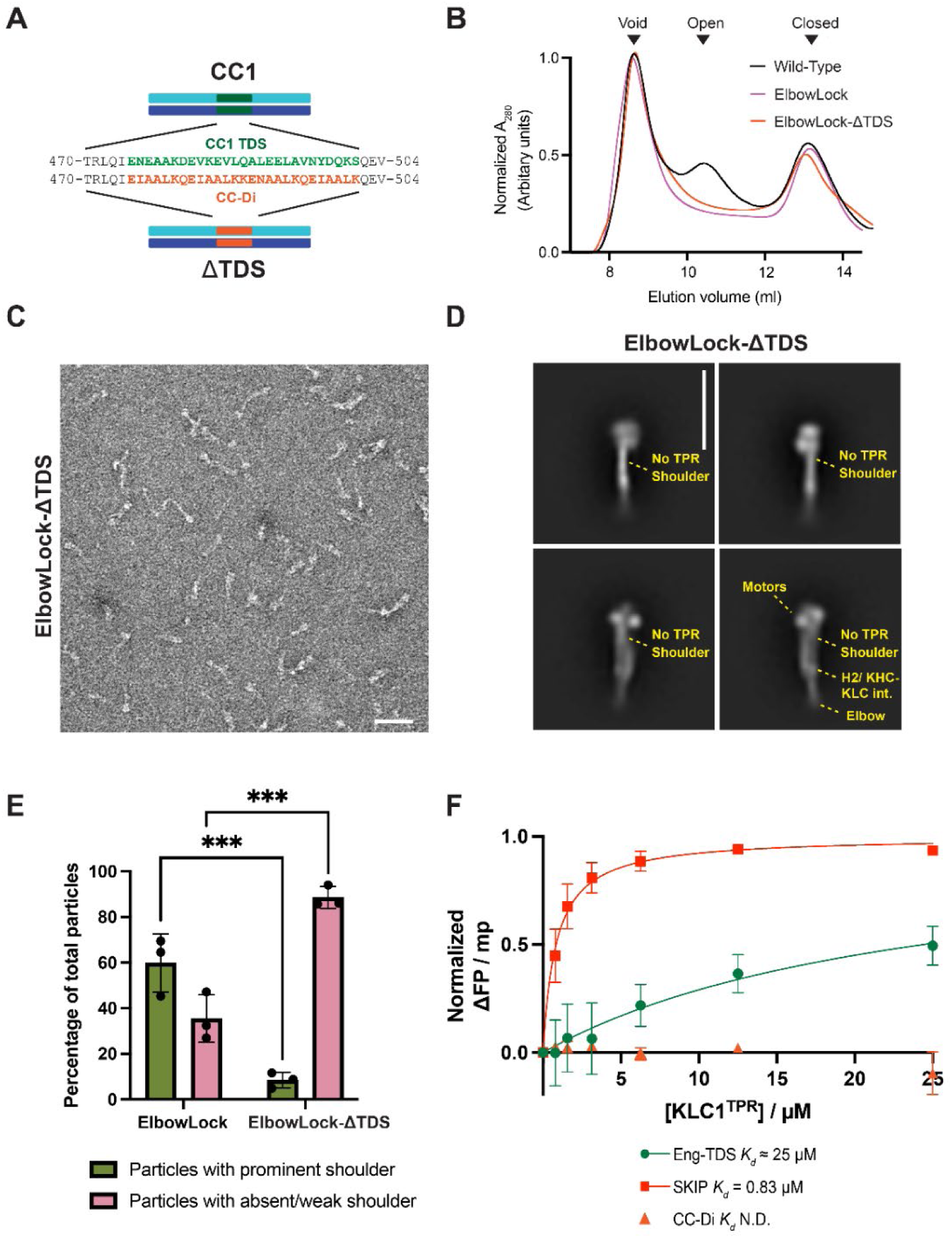
The CC1 TDS is required for the formation of the kinesin-1 shoulder. (A) Schematic showing strategy to delete the TDS in CC1 and replace it with an equivalent length of a *de novo* designed homodimeric coiled coil, CC-Di. (B) SEC traces showing the elution profile of wild type, ElbowLock, and ElbowLock-ΔTDS proteins. Constructs in the ElbowLock background elute exclusively in the peak associated with closed lambda particles. Representative of 3 independent experiments. (C) Representative NS electron micrographs (from 3 independent experiments) showing ElbowLock and ElbowLock-ΔTDS complexes. Scale bar is 50 nm (D) Selected 2D classes from NS-EM highlighting the presence of the shoulder in ElbowLock but not ElbowLock-ΔTDS complexes. The scale bar is 26 nm. Full sets of classes are provided in Fig. S4 (E) Quantification of number of particles in classes with and without a prominent shoulder of 3 independent experiments. *** indicates p<0.001 (F) Fluorescence polarisation binding assays show fluorescently labelled TDS and SKIP (W-acidic motif, positive control) peptides, but not the control peptide CC-Di, bind to isolated TPR domains. Error bars show S.E.M. from 3 replicates.

### The TDS binds directly to the KLC TPR domain

To determine whether the TPR binds TDS directly, we measured the interaction between fluorescently labelled synthetic peptides corresponding to the TDS region and the TPR domain directly using fluorescence polarisation (FP) assays. Short peptides from the TDS region were unfolded in the solution, which we attributed to an imperfect hydrophobic repeat pattern, lacking leucine residues typical of the cores of dimeric coiled coils. Therefore, we made 3 mutations of core residues (A479, V486, K500) to leucine and templated folding using 7 flanking residues on either side of the sequence from CC-Di **(Fig. S6A)**. Circular dichroism (CD) spectroscopy and analytical ultracentrifugation (AUC) confirmed that the resulting peptide (Eng-TDS) formed a helical dimer **(Fig. S6B, C, D, E)**. Subsequent FP assays showed that isolated KLC1-TPR interacted with Eng-TDS with low affinity (approx. *Kd* of 25 µM), but not with CC-Di alone. **(Fig. 2F)**. We obtained similar results for KLC2-TPR with tighter binding to Eng-TDS **(Fig. S6F)** (*Kd* = 2.7 µM), suggesting that this interaction is conserved between KLC paralogues. We note that these binding measurements are for an inter-molecular process but in the context of the holoenzyme the interaction is effectively intra-molecular because the TPR domain is tethered to the coiled-coil scaffold. Therefore, the effective local concentration is substantially higher, resulting in a high equilibrium fraction of bound complexes, consistent with our EM experiments. Prior work shows that constructs with deletions of the first helix of TPR1 of KLC2 result in unfolding of TPR1 (*35, 52*). Therefore, to test a key role for the TPR1 interface, we deleted the first helix of TPR1. This effectively abrogated any detectable binding **(Fig. S6G)**. Together, this modelling, electron microscopy analysis and biophysical experiments clearly demonstrate that the TDS provides a TPR domain docking site on the lambda particle.

### Cargo adaptor binding dislocates the TPR shoulder

To understand the consequences of cargo binding, we incorporated an activating W-/Y-acidic cargo adaptor ligand into the complex (KinTag) via fusion on a flexible linker at the carboxy-terminus of KLC; i.e., an intramolecular mimic of stable cargo binding, following a similar strategy used to solve the structure by X-ray crystallography (*36*) **(Fig. 3A)**. In a wild-type background, this resulted in a shift in the SEC elution profile from the two peaks corresponding to open and closed conformations to a single peak at an intermediate position **(Fig. 3B)**, which negative stain EM analysis revealed consisted of a mixture of open and closed particles, but with no discrete intermediate state **(Fig. S7)**. From this, we conclude that the main effect of SLiM-TPR interaction is to increase the rate of exchange between open and closed conformations such that they are no longer resolved by SEC. Supporting this, additional incorporation of the closed-stabilising ElbowLock modification shifted the equilibrium to a position corresponding to the closed peak, demonstrating that the cargo-induced SEC shift can be supressed by restricting conformational flexibility at the elbow, and therefore results from opening the complex **(Fig. 3B)**.

**Figure 3.**
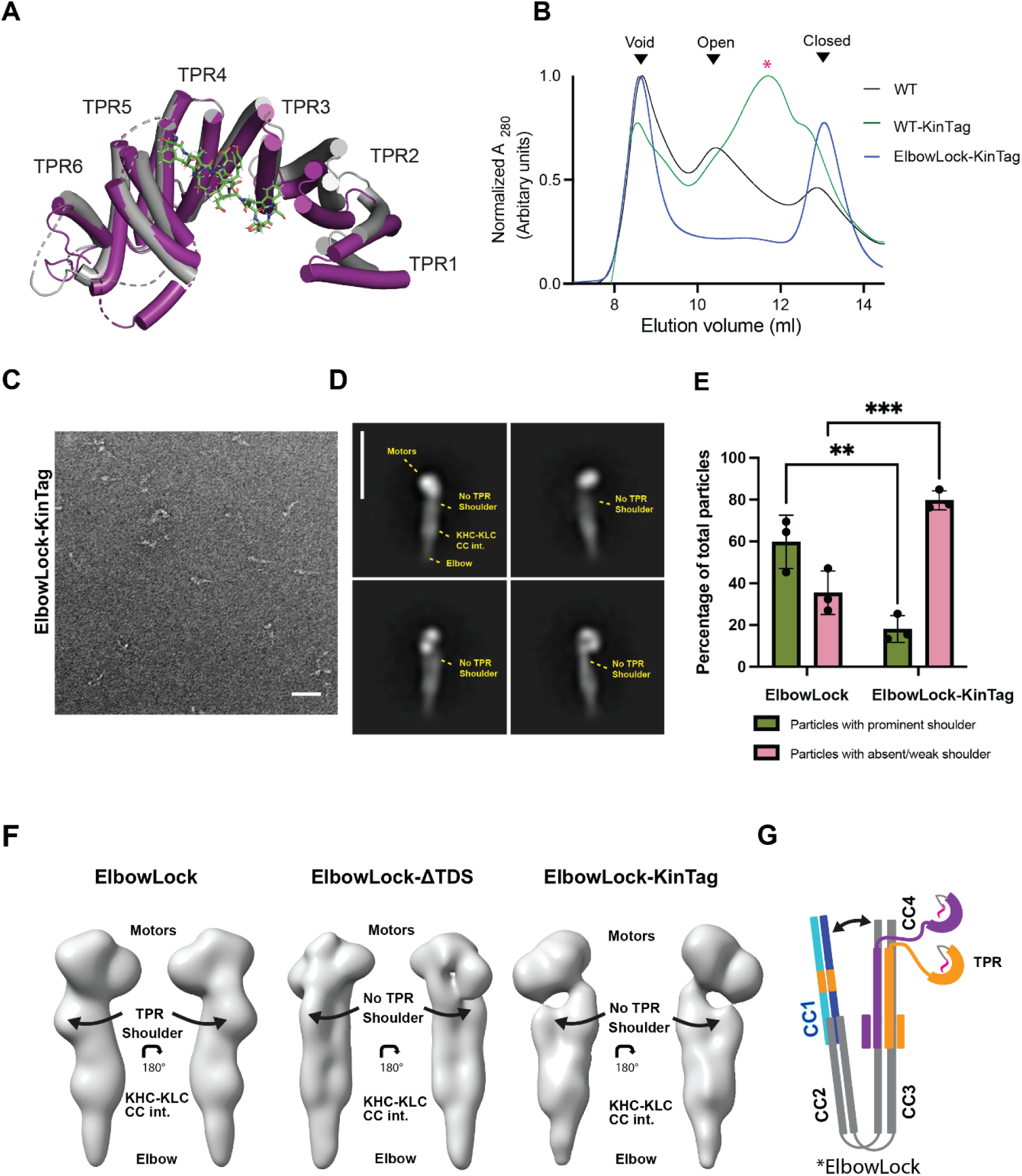
Adaptor-binding to KLC-TPR dislocates the kinesin-1 shoulder. (A) Comparison of X-ray crystal structures of ligand-free (grey, pdb: 3nf1) and KinTag (W/Y acidic)-ligand bound (purple, pdb: 6swu) TPR domains highlighting the ligand-binding site and ligand-induced change in TPR curvature. (B) SEC traces showing elution profile of wild type, ElbowLock and ElbowLock-KinTag proteins. (C) Representative NS electron micrographs (from 3 independent experiments) showing ElbowLock and ElbowLock-KinTag complexes. Scale bar = 50 nm. (D) 2D classes from NS-EM highlighting the loss of the shoulder in ElbowLock-KinTag complexes. The scale bar is 26 nm. (E) Quantification of the percentage of particles in classes with and without a prominent shoulder from 3 independent experiments. ** indicates p 0.001 and *** indicates p < 0.001 (F) 3D-reconstructions from the ElbowLock, ElbowLock-ΔTDS and ElbowLock-KinTag datasets show dislocation of the shoulder either upon deletion of the TDS or incorporation of KinTag. (G) Model showing dislocation of the shoulder induced upon KinTag binding.

Analysis of the resulting ElbowLock-KinTag particles by NS-EM and 2D classification resulted in clear loss of the TPR shoulder (**Fig. 3C-E)**, showing that the intramolecular adaptor interaction promotes TPR dissociation from the closed complex. To substantiate this further, we generated 3D models of ElbowLock, ElbowLock-ΔTDS and ElbowLock-KinTag complexes from the NS-EM data **(Fig. S8)**. For ElbowLock complexes, this resulted in classes with and without a prominent shoulder, in agreement with 2D classification. For ElbowLock-ΔTDS and ElbowLock-KinTag complexes, no prominent shoulder containing classes were observed. Moreover, these modifications appeared to result in a slight loosening of the whole coiled-coil assembly **(Fig. 3F)**; note the indentations between the densities of the two main coiled-coil regions, which we interpret as the start of a separation of these two domains that is restricted by the ElbowLock mutation. Consistent with a KinTag-induced inhibition of TDS-TPR association, inclusion of KinTag on the KLC-TPR carboxy-terminus effectively abrogated any detectable binding to Eng-TDS, or its direct competitor, the W-acidic motif of SKIP, in FP assays **(Fig. S6H)**. Together, these data support a model where adaptor-TPR interaction dislocates the TPR shoulder from the complex **(Fig. 3G)**.

### Cargo-mediated dislocation of the shoulder enhances motor domain accessibility

To determine how the local structural changes from adaptor binding and shoulder dislocation affected the dynamics of kinesin-1 complexes in solution, as directly and least invasively as possible, and without the risk of cross-linker artefacts, we recorded millisecond time-resolved hydrogen/deuterium-exchange mass spectrometry (HDX-MS) time courses, resolved at the peptide level **(Fig. S9, S10, Table S1)** (*53–57*). Sequence coverage was good (overall 88%) with the exception of KHC-CC0 (neck coil) and the acidic-linker region that connects the KLC coiled-coil to the TPR domains where coverage was more limited.

To begin, we established baseline parameters by comparing the wild-type and DeltaElbow complexes. The latter stabilizes the fully extended conformation by promoting helical readthrough at the elbow and thus forces TPR undocking from CC1 (*32, 34*). The data for wild type were subtracted from those of the DeltaElbow to generate difference plots **(Fig. S11A, S12A)**, which were then mapped onto the X-ray crystal structures of the motor and TPR domains **(Fig. 4)** (*52, 58*).

**Figure 4.**
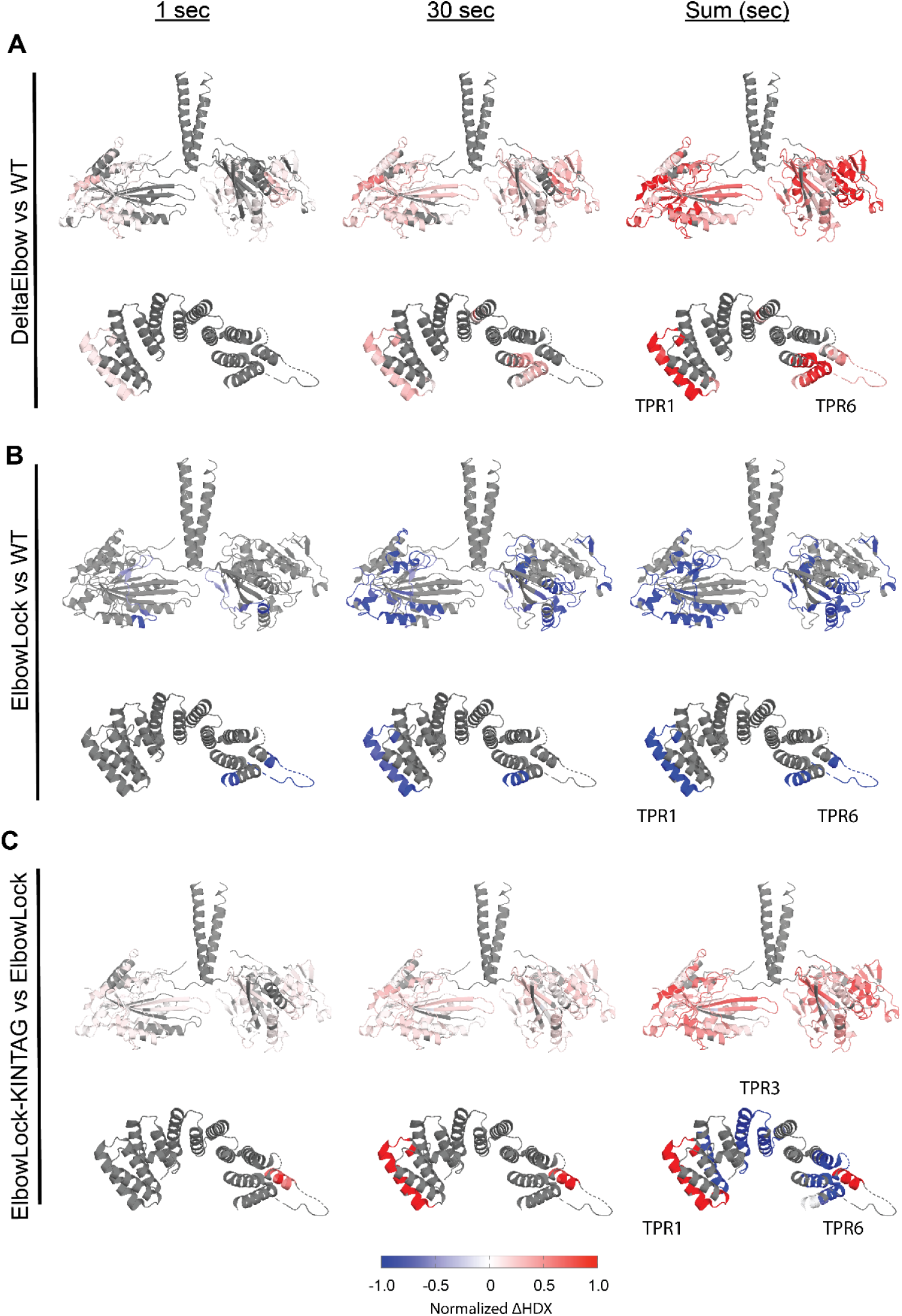
Adaptor-binding to the KLC TPR domain promotes motor accessibility. Summaries of comparative HDX-MS analysis of wild type, DeltaElbow, ElbowLock and ElbowLock-Kintag complexes. At each time point, the rates of exchange were subtracted as indicated (X minus Y) and filtered and filtered using a hybrid significance test (global significance and Welch’s t-test, then transposed onto X-ray crystal structures of the motor domains (pdb: 3kin) and TPR domain (pdb: 3nf1). Grey indicates non-significant differences or limited peptide coverage. (A) Comparison of deltaElbow (X) against wild type (Y). (B) ElbowLock (X) against wild type (Y). (C) ElbowLock-Kintag (X) against ElbowLock (Y). Relative protection (blue) indicates less solvent exposure or increased H-bonding for X, while deprotection (red) indicates more solvent exposure or H-bond loss for X.

First, as a general observation for the KHC chain, the largest differences in HDX were observed outside of its coiled-coil (CC) stalk. This is consistent with hydrogen-bonded α-helical structure and the known stabilities of coiled-coil assemblies in general that is protective against hydrogen-exchange and may partially mask changes in solvent accessibility (*59*). Nonetheless, there were several small but statistically significant differences in labelling between the two constructs in the KHC CCs consistent with the opening of the DeltaElbow complex. These included deprotection/increased accessibility at the proposed motor-CC1 interface (*22, 33*), the TDS, and the KHC-KLC interface in CC3, running into the amino terminus of CC4 **(Fig. S11A)**. Interestingly, for the KLC chain in the open state, we observed more deprotection/increased accessibility in its CC and even protection at its amino terminus suggesting larger-scale activation associated structural changes in this component of the kinesin complex **(Fig. S12A)**.

Second and strikingly, the DeltaElbow modification significantly increased HDX rates of the motor domains **(Fig. 4A)**. This is consistent with their liberation from the folded-back and locked conformation of the lambda particle (*22*). The increase in HDX was not tightly localised. Rather, the motor domain as a whole was significantly deprotected. This is consistent with enhanced solvent accessibility and increased local subdomain dynamics throughout the motors in response to opening of the elbow. In addition, opening the complex resulted in significant deprotection/increased solvent accessibility of TPR1. This is consistent with its role in docking the TPR onto the CC assembly and its forced undocking by helical readthrough in the DeltaElbow variant. We also observed additional deprotection of TPR6, extending through to the C-terminal sequence that follows the TPR, indicating possible occlusion of these features in the inhibited state.

Next, we compared the ElbowLock (stabilised closed) and wild-type complexes. Opposite to DeltaElbow, both the motor domains and the TPR1 sites were more protected/less accessible in ElbowLock variant **(Fig. 4B, S11B, S12B)**. There was also protection of the C-terminal tail of KHC. Thus, shifting the open-closed equilibrium to either of its extremes enables visualization of the accessibility of key interfaces in the complex that maintain autoinhibition, and support our findings emerging from AF3 modelling, negative-stain EM, and biophysical analyses.

Finally, we compared the ElbowLock-KinTag and ElbowLock complexes to examine the consequences of adaptor binding in the closed state **(Fig. 4C, S11C, S12C)**. As expected, inclusion of KinTag significantly decreased HDX in, and therefore stabilized, the central TPR repeats at its structurally characterised binding site (*36*). This was juxtaposed with the striking deprotection of TPR1, which is consistent with ligand-induced TPR domain undocking from the lambda particle at this epitope **(Fig. 3)**. The motor domains were also deprotected. Changes on the KHC CC were again much lower in magnitude, but we did note statistically significant deprotection at the predicted motor-CC1 docking site and C-terminal tail, suggesting that TPR-adaptor binding triggers release of the motors from their folded back, locked conformation.

Together, these data provide direct in-solution measurements revealing both predicted and unanticipated changes in solvent accessibility associated with the opening and closing of the complex that demonstrate allosteric coupling between adaptor-induced dislocation of the TPR shoulder and increased motor domain accessibility.

### Opening the kinesin-1 complex enhances its association with MAP7

Thus far, our findings support a model where cargo-binding destabilises the compact conformation and so, may unmask protein-protein interaction interfaces that facilitate subsequent steps in the activation pathway. The microtubule associated protein MAP7 is an essential kinesin-1 cofactor that promotes binding to microtubules and activation. Its binding site has been biochemically mapped to a short region of CC1 that overlaps the TDS (*60–67*). Indeed, AlphaFold3 predictions suggest that although the TPRs and MAP7 may bind to CC1 via distinct mechanisms (near perpendicular TPR helix-turn-helix motif vs parallel single MAP7 alpha helix), their binding sites directly overlap, and as such, they may compete for KHC binding **(Fig. 5A)**. One prediction of this model is that opening of the complex at the Elbow to force TPR undocking would enhance binding to MAP7. To test this, we immunoprecipitated GFP-tagged MAP7 from cells expressing wildtype, DeltaElbow and ElbowLock kinesin-1 complexes. This resulted in a roughly 5-fold increase in kinesin-1 associated with MAP7 in the DeltaElbow background, compared to wildtype or ElbowLock constructs **(Fig 5B,C)**. We also noted slightly elevated expression of ElbowLock complexes and slightly lower expression of DeltaElbow complexes, suggesting that opening/closing of the complex could impact on kinesin-1 turnover. Similarly, in immunofluorescence experiments, the DeltaElbow modification resulted in a striking increase in kinesin-1 associated with MAP7-positive microtubules and pronounced microtubule bundling **(Fig 5D)**. Together, these data reveal a coupling between kinesin-1 conformational state and its capacity to associate with the microtubule associated co-factor MAP7.

**Figure 5.**
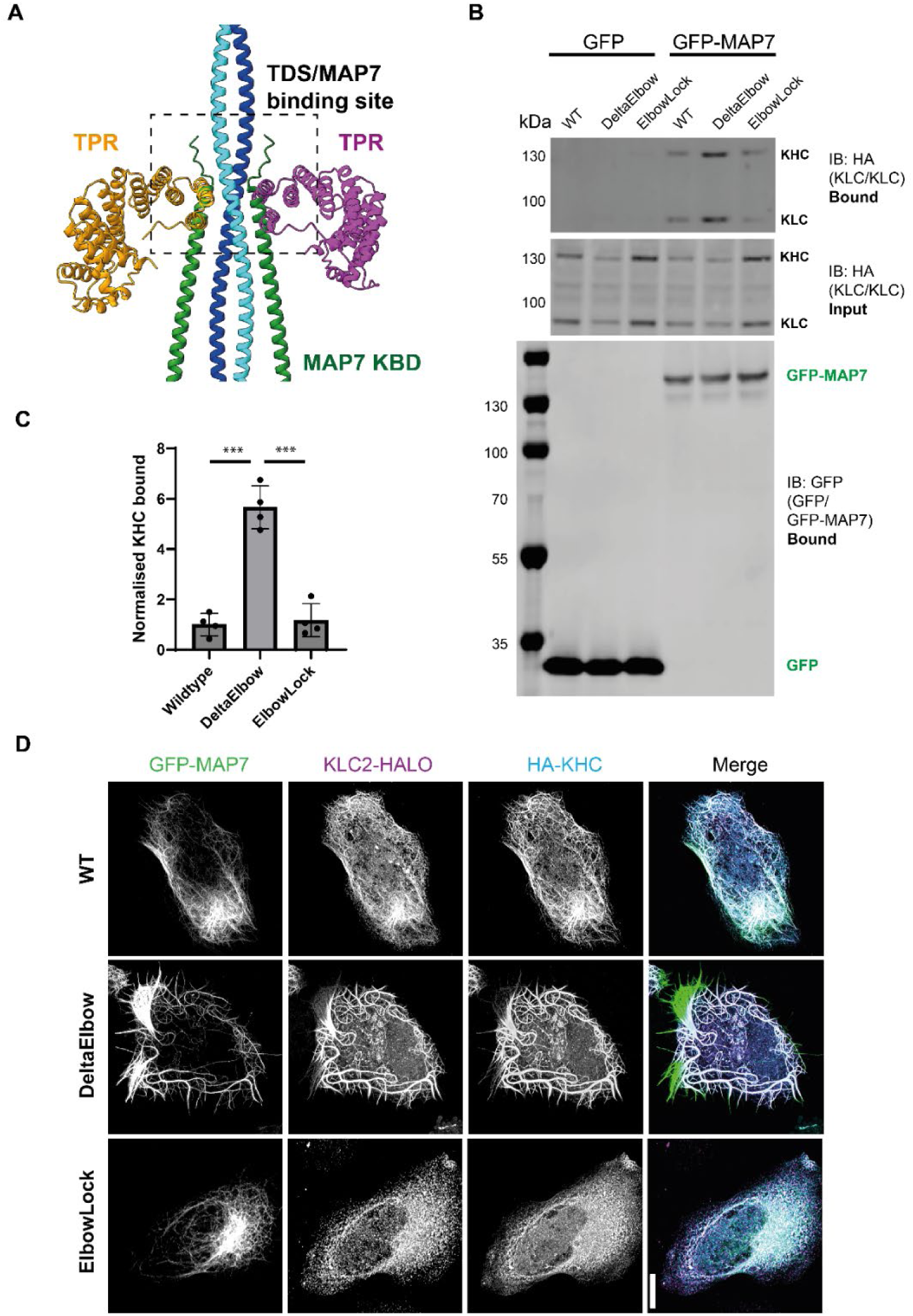
Opening the kinesin-1 complex promotes its association with MAP7. (A) Overlay of two Alphafold3 models of a tetrameric CC1/TPR assembly and a tetrameric CC1/MAP7-kinesin binding domain assembly, aligned on CC1. For clarity, only one CC1 model is shown. (B) GFP-MAP7 immunoprecipitation experiments showing enhanced association of KHC and KLC in the DeltaElbow background. Cells were transfected with the indicated constructs (GFP/GFP-MAP7, HA-KLC2, HA-KIF5C (WT, ElbowLock, DeltaElbow). Complexes were immunoprecipitated using GFP-TRAP beads, and input and bound samples were analysed using western blotting with anti-GFP and anti-HA antibodies. (C) Quantification of b. from 4 independent experiments. Error bars show S.E.M. ***=p<0.001 using 1-way ANOVA with Tukey’s multiple comparison test. (D) Immunofluorescence images of HeLa cells transfected with GFP-MAP7, HA-KHC, and KLC2-Halo (TMR-labelled) showing enhance association of kinesin-1 with GFP-MAP7 positive microtubules in the DeltaElbow background. Images are representative of 3 independent experiments. Scale bar is 20 µm.

## Discussion

Over the past two decades, studies have identified a key role for SLiM-containing adaptor proteins in recruiting and activating kinesin-1 by binding to the KLC-TPR domains (*2, 26, 35, 36, 39, 40, 42–45*). Recent work has revealed the low-resolution architecture of the autoinhibited lambda state, including the TPR shoulder resting on the coiled-coil body (*22, 32, 33*). Increasing evidence suggests that a balanced equilibrium between inhibited and active states ensures responsiveness to stimuli (*34, 40, 68*). The challenge is to understand how adaptor binding to the TPR drives the transition from the closed, inhibited state to the open, active state; intrinsic conformational dynamics and unitarily low-affinity, cooperative, motor-cargo interactions obscure key intermediates in this pathway. To address this here, we have used *de novo* protein design and protein engineering to stabilise these intermediates alongside biophysical, biochemical, and electron microscopy measurements of the complete heterotetrametric holoenzyme. Our results lead us to propose a model where adaptor binding dislocates the TPR shoulder, destabilizing the autoinhibited state, enhancing motor domain accessibility and promoting the transition from the inactive closed state to the open forms **(Fig. 6)**.

**Figure 6.**
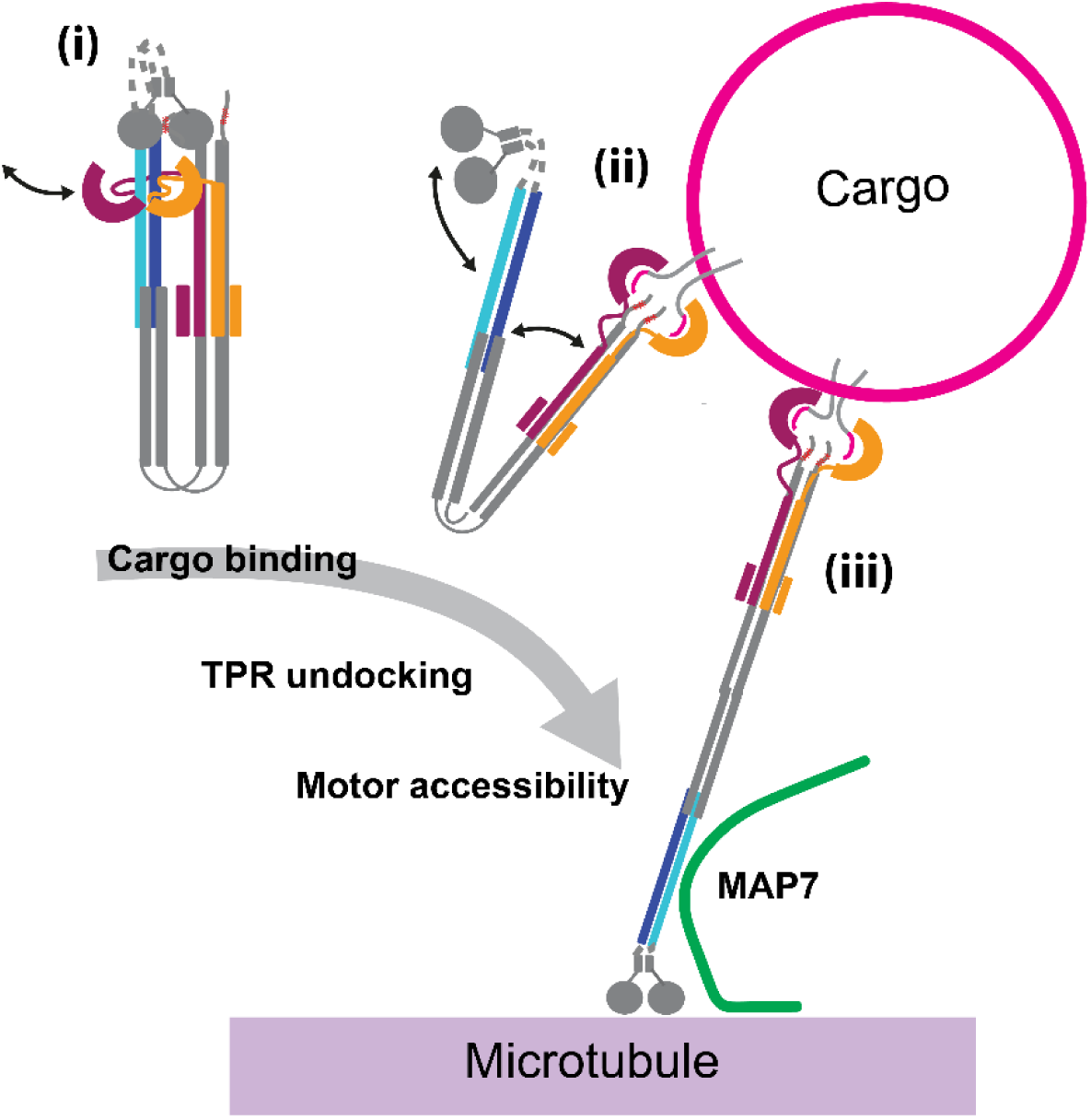
Model for cargo-mediated initiation of kinesin-1 activation. In the autoinhibited state, a TPR domain(s) are docked at the TDS forming the shoulder. i. Binding to the cargo adaptor dislocates the TPR shoulder and promotes motor domain accessibly, and initiates separation of the coiled-coil domains (ii). This allosteric coupling between cargo and motor would facilitate recruitment to microtubules (supported by MAP7) and subsequent transport (iii).

Integration of the present findings with prior work suggests a unified model for the role of TPR binding cargo adaptors in motor activation. We reveal that TPR1, the first TPR repeat, forms a key interface for TPR domain-CC association. TPR1 also binds the adaptor protein JIP3 via its coiled-coil leucine zipper, resembling TDS interaction predictions. However, a detailed evolutionary analysis in the same study suggests that TPR1 plays a distinct role beyond cargo binding (*50*). Moreover, a comparison of high-resolution X-ray crystal structures of KLC-TPR domains, with and without SLiM cargo, demonstrates structural plasticity in TPR1, which may enable ligand binding to modulate TPR1 orientation alongside larger scale conformational changes resulting from an increase in TPR domain curvature in response to SLiM binding (*24, 35, 45, 69*). We propose that SLiM ligand binding to the TPR groove drives conformational changes that reduces TPR1 affinity for the TDS to dislocate the shoulder; JIP3 may compete for the same site on TPR1 to the same effect. Our previous work has shown that SLiM-TPR interaction gates access to important cargo and microtubule binding sites on CC4, that is predicted to be closely juxtaposed to CC1 in models of the inhibited state (*24, 37, 47*). Our computational predictions do not indicate a direct interaction between KLC TPR and CC4 although they are predicted to be in proximity. Whilst we do not exclude this possibility of a direct interaction, we favour a model where removal of steric blocking effects from the bulky TPRs and docked motors on CC1, as well as opening of the coiled-coil assembly enables subsequent steps such as cargo binding to additional sites on CC4.

The TDS is located within biochemically defined binding site for the essential kinesin-1 cofactor MAP7 (*60–67*). Recent studies show clear cooperativity between MAP7 and W-acidic SLiM adaptors in motor activation (*40, 70*). This work provides a compelling explanation for this co-operativity: shoulder dislocation would be predicted to destabilise autoinhibition and expose the MAP7 binding site, facilitating subsequent activation steps such as enhanced microtubule recruitment and transition to and/or maintenance of the fully extended state **(Fig. 6)**. Consistent with this, our models suggest that CC1 binding to MAP7 or KLC TPR would be mutually exclusive *if* the symmetrical binding sites on CC1 were both occupied (2xMAP7 or 2xKLC-TPR). However, it is not clear at this point whether this is the case. In the absence of nanobody-mediated TPR stabilisation, negative stain EM hints at an asymmetric TPR conformation and we remain open to the possibility that only one TPR is bound at one time and that they may exchange. This would, in principle, leave one CC1 binding site free to interact with MAP7 and one TPR domain free to make first contact with cargo.

It will be important to understand this dynamic interplay and to sequence these events. A cargo initiated opening up of the complex is consistent a model for MAP7 function where it facilitates tethered diffusion to bypass obstacles on the microtubule and enhances kinesin-1 processivity (*61, 62*), and activation without cargo-recognition would seem counter-productive. However, recently, MAP7 binding to KHC was shown to promote KLC binding to W-acidic and Y-acidic cargo (*70*). Here, we have used a high-affinity ligand to mimic stable cargo attachment, but natural motifs typically bind with low affinity and act cooperatively with additional protein-protein and protein-lipid interactions (*27, 35, 39, 47, 71*). Therefore, one might also consider a two-way switch model where MAP7 binding also facilitates shoulder dislocation to modulate cargo binding properties of the KLC TPR domains. In either case, this points to a critical interplay between regulator and cargo, converging on CC1, establishing a new target for intervention in kinesin-1 mediated transport processes, and design of new reagents to target this site will likely be critical to dissect its dynamic functions.

In summary, our findings demonstrate that kinesin-1 autoinhibition and initiation of activation by SLiM-cargo pivots on the association and dislocation of its TPR shoulder from the coiled-coil scaffold. One important implication of these findings is that kinesin-1 is unlikely to be stably inhibited when bound to cargo but instead may be primed for further regulatory inputs that control its activity. This may be important for the coordination of bidirectional transport in multi-motor/adaptor systems.

## Acknowledgements

This work was funded by BBSRC grants BB/W005581/1 and BB/Z517276/1. JAC was supported by an EPSRC-funded Doctoral Prize Fellowship. MPD acknowledges support from a Lister Institute of Preventative Medicine Fellowship. CS acknowledges funding by a Wellcome Trust Investigator award (210701/Z/18/Z). JJP and MK acknowledge funding from UKRI Future Leaders Fellowship (MR/T02223X/1). We thank the University of Bristol School of Chemistry Mass Spectrometry Facility for access to the EPSRC-funded Bruker Ultraflex MALDI-TOF instrument (EP/K03927X/1), the BBSRC-funded BrisSynBio centre for access to peptide synthesis and the plate reader (BB/L01386X/1) and the Wolfson Bioimaging Facility for access to this Talos L120C microscope supported by BBSRC equipment grant (BB/X019799/1). We are grateful for the assistance and access to equipment at the GW4 Facility for High-Resolution Electron Cryo-Microscopy, funded by the Wellcome Trust (202904/Z/16/Z and 206181/Z/17/Z). We are grateful to Alex Walker and Bram Mylemans for helpful discussions and Ferdos Abid Ali for comments on the manuscript.

## Author Contributions

SS purified kinesin-1 complexes, performed SEC, and NS-EM experiments and analysis. JAC designed and engineered the ΔTDS KIF5C construct, designed, synthesised and characterised peptides for FP binding assays, purified TPR proteins and performed and analysed FP experiments. MK, SS and JJP designed, carried out and analysed HDX-MS experiments. SS, SKNY, CS and MPD analysed cryoEM data. JW designed and engineered the KLC1-KinTag fusion protein and other constructs. UB and JM supported NS and cryo-EM sample preparation and data acquisition. XL performed AUC assays. MPD, DNW and SS wrote the paper with input from all the authors.

## Materials and Methods

### AlphaFold3 modelling

All of the models shown in the manuscript were predicted using the Alphafold3 server (*72*). The algorithm was asked to model a heterotetramer composed of KIF5C (NP_001101200.1) and KLC1A (NP_032476.2) with varying input sequences described in the figure legends. Predicted aligned error statistics are as output by the software with manual annotations to highlight important features. For the models presented in Fig. 5, the algorithm was asked to model a heterotetramer of KIF5C CC1 (S410-G558) with the kinesin binding helix of MAP7 (NP_001185564.1, P467-K610), or KLC1 TPR domain (NP_032476.2, G205-K495). Similar models and predicted binding sites were obtained for heterotrimers (2xCC1, 1xMAP7/TPR). Output models were visualised/aligned and prepared for publication using in UCSF Chimera (*73*).

### Constructs for protein expression

A 2-plasmid system to express heterotetrameric kinesin-1 in *E. coli* cells has been described previously (*32, 34*) and constructs used in this study were derived from these. Wild-type kinesin heavy chain was KIF5C expressed in the pMW bacterial expression vector with an N-terminal 6X His tag. KLC1 was expressed pET28a with the His tag removed by site-directed mutagenesis. ElbowLock was originally designed by deleting five amino acids from the elbow loop (E685-L690) (*34*), and ElbowLock-ΔTDS was prepared by additionally replacing E475-S501) in CC1 with 4-heptads of CC-Di through subcloning of a synthetic gene fragment. KLC1-KinTag construct was prepared by fusing KinTag peptide via a (Thr-Gly-Ser)_10_-Gly flexible linker at C-terminal of KLC1 (*36*). The TPR domain of KLC1 (A211-A495) and KLC2 (A196-K480, delta helix 1 P219-K480) were cloned in the bacterial expression vector pET28His-Thrombin and have been described previously (*24, 35*). Anti-TPR nanobody (sequence from pdb: 6fv0)(*45*) was codon optimised and synthesised and cloned into NcoI and XhoI site of pet22b with N-terminal pelB sequence and C-terminal 6X His tag, by Genscript Inc. The identity of all plasmids was confirmed by DNA sequencing.

### Protein expression and purification

All kinesin-1 constructs were expressed in *E. coli* BL21(DE3) cells as a heterotetramer adapting the two-plasmid system as described previously (*32, 34*). Briefly, expression plasmids encoding individual KIF5C constructs and KLC1 constructs were transformed into *E. coli* BL21(DE3). Positive clones were selected based on a double antibiotic selection marker. Cells were cultured in LB medium (miller) supplemented with ampicillin (50 µg/ml) and kanamycin (50 µg/ml) in a shaking incubator under conditions of 37°C/180 rpm. When the optical density (OD 600) measured at 600 nm reached 0.6, the temperature was reduced to 18°C. Protein expression was induced by addition of 0.2 µM IPTG. Cultures were incubated overnight with shaking at 18°C. Cells were harvested by centrifugation at 5000 g for 15 min at 4°C. Cells were resuspended in Buffer A (40 mM Hepes (pH 7.4), 500 mM NaCl, 40 mM imidazole, 5% glycerol, and 5 mM β-mercaptoethanol), supplemented with protease inhibitors. Lysis was carried out by sonication (70% amplitude, 5s on/15 s off) for 7.5 min on ice. Lysates were centrifuged at 40,000g for 45 min at 4°C to obtain supernatant and filtered through a 0.45 µm membrane. The supernatant was loaded onto a His-Trap (1ml) column equilibrated with Buffer A. The column was washed with at least 50 CV of Buffer A with 60 mM Imidazole. Bound proteins were eluted using a gradient of Buffer A with increasing concentrations up to 500 mM of Imidazole. Eluted protein fractions were analysed on SDS-PAGE, then concentrated using a 30,000-Da MWCO ultrafiltration device up to 500 µl and immediately loaded on a size exclusion chromatography (SEC) column. SEC was carried out using a Superose 6 10/300 column pre-equilibrated with 20 mM Hepes (pH 7.4), 150 mM NaCl, 1 mM MgCl2, 0.1 mM ADP, and 0.5 mM TCEP. Eluted protein fractions were analysed using SDS-PAGE. Appropriate fractions were aliquoted and stored at -80°C freezer and used further for downstream applications.

TPR constructs were expressed in *E. coli* BL21(DE3) and affinity purified in buffer (25 mM Hepes pH 7.4, 500 mM NaCl, 5 mM β-mercaptoethanol) over a gradient of 20 mM to 500 mM imidazole on a His-Trap (1ml) column (*24, 35*). SEC was carried out using a HiLoad 16/600 Superdex 75 prep grade column (GE Healthcare) equilibrated with 25 mM HEPES (pH 7.4), 500 mM NaCl, 5 mM β-mercaptoethanol. Eluted fractions were analysed on SDS-PAGE.

Nanobody was expressed in *E. coli* BL21(DE3) and isolated form periplasmic space in 0.2 M Tris pH 8, 0.5 M EDTA, 0.5 M Sucrose and purified on His-Trap (1ml) column. SEC was carried out using Superose 6 10/300 column pre-equilibrated with PBS. Eluted fractions were concentrated and mixed in 10-fold excess with SEC purified ElbowLock (5.56 µM: 66 µM). Eluted complexes were analysed on SDS-PAGE.

### Negative stain electron microscopy

Protein aliquots (50 µl) were thawed from −80°C storage and cross-linked with 0.6 µl of 50 mM BS3 (final concentration 0.59 mM) at room temperature for 30 minutes. After cross-linking, the protein sample was placed on ice, and serial dilutions were prepared to achieve optimal single-particle distribution on grids. Formvar/carbon-coated 300-mesh copper grids were glow-discharged for 30 seconds at 20 mA. A 5 µl aliquot of the protein sample was applied to the grid and incubated for 1 minute before blotting with filter paper. The grid was subsequently stained by sequentially picking and blotting into three 5 µl drops of 3% uranyl acetate. The first drop served as a quick rinse, the second was left on the grid for 2 minutes, and the third also served as a quick rinse. The prepared grids were air-dried and subsequently imaged on a Talos L120C transmission electron microscope operated at 120 kV, equipped with a 4k × 4k Ceta CMOS camera. Micrographs were acquired at a magnification of ×57,000, corresponding to a pixel size of 2.48 Å/pixel. Datasets of approximately 500 micrographs were acquired using EPU automated collection software for all samples. The total electron dose was 52 e−/Å², and micrographs were recorded with defocus values ranging from −1.5 µm to −2.5 µm.

### Negative stain electron microscopy image processing

Micrographs were analysed using CryoSPARC (*74*). After importing the micrographs, CTF correction was performed using the Patch CTF tool. Approximately 1,000 particles were manually picked and subjected to 2D classification to generate templates for automated particle picking. Using these templates, particles were autopicked, extracted (63.4 nm box size), and fourier-cropped to a pixel size of 4.96 Å/pixel. Extracted particles were 2D classified into 200 classes. A cleanup process was conducted by discarding particles that did not resemble kinesins through iterative rounds of curation and 2D classification. The number of particles in classes showing full ‘side views’ of the complex were counted and distinguished as particles with or without a prominent shoulder. The data quantified in this way was statistically analysed using 2-way ANOVA (using Sidak’s multiple comparison test). For figure presentation only, and to allow for intuitive comparison to AlphaFold models, final particle sets were reclassified with the CryoSPARC ‘align filament classes vertically’ option ticked. This resulted in no obvious morphological differences, but produced classes aligned on long axis of the complex. For 3D reconstruction, cleaned particle sets were used to generate four ab-initio models in CryoSPARC. A heterogeneous refinement of ab-initio models was performed, followed by homogeneous refinement, with alignment resolution restricted to 20 Å. Resulting models were low-pass filtered to 40 Å resolution and visualized using ChimeraX (*73*).

### Cryo-electron microscopy and image processing

Purified cross-linked ElbowLock complexes (5 μl at 3 mg/ml) were applied to glow-discharged Quantifoil R1.2/1.3 200-mesh Cu grids. The grids were blotted for 2 seconds at 4 °C and 100% humidity, then flash-frozen in 37% ethane/propane mix kept at approximately at –195°C using an FEI Vitrobot Mark IV (Thermo Fisher). Images were acquired on a Talos Artica microscope (FEI) operating at 200 kV with a Gatan K2 Summit direct detector. Automatic image collection was performed using EPU software (Thermo Fisher). A total of 3943 micrographs were captured with a total dose of 58 e/Å², dose-fractionated into 64 movie frames at a defocus ranging from −1 µm to −2.5 µm. Images were recorded at 130,000× nominal magnification, resulting in a pixel size of 1.05 Å per pixel. Data processing was carried out in CryoSPARC (*74*). Motion correction and CTF estimation were performed using Patch Motion Correction and Patch CTF, respectively. Particles (focussing on full length side views) were manually picked from selected micrographs to create templates for automated particle picking. The autopicked particles were extracted and binned to 4.2 Å/pixel with a box size of 128 pixels / 53.76 nm Extracted particles underwent multiple rounds of 2D classification to produce the class averages shown in Fig. 1. For comparison to the AF3 model, simulated density was generated using the molmap command in ChimeraX (*73*) filtering to 15 Å. and projections were generated/selected automatically using the Reference Based Auto Selected 2D function in CryoSPARC.

### Hydrogen-deuterium exchange mass spectrometry

Hydrogen-deuterium exchange (HDX) was conducted using “ms2min”, a fully automated, millisecond-resolution HDX labelling online quench-flow device (Applied Photophysics Ltd), which was connected to a Waters HDX manager. In the labelling experiments, 14 µL (10 µM) of DeltaElbow, wildtype, ElbowLock and ElbowLock-KinTag was introduced into the labelling mixer separately. The HDX reaction was initiated by diluting the sample 20-fold with a labelling buffer at 20°C. This buffer consisted of 20 mM HEPES, 150 mM NaCl, 1 mM MgCl2, 0.1 mM ADP, 0.5 mM TCEP in D2O, adjusted to a pH of 7.40 at 20°C. Samples were labelled at seven different time points (0.3, 0.5, 1, 3, 10, 30, 300) in triplicate. The HDX reaction was immediately quenched by mixing 1:1 with quench buffer (8 M urea, pH 2.55, at 0°C), and the sample was digested online using a Waters Enzymate pepsin column. The resulting peptides were trapped on a 2.1 × 5 mm VanGuard ACQUITY BEH C18 column (Waters) for 3 minutes at a flow rate of 155 µL/min and then separated on a 1 × 100 mm ACQUITY BEH C18 column (1.7 μm particle size) using a 7-minute linear gradient of 5-40% acetonitrile with 0.1% formic acid. Peptides were analysed using a Synapt G2-Si mass spectrometer (Waters) in HDMSE mode across a mass range of 50–2000 m/z. Instrument settings were as follows: capillary voltage of 3.0 kV, cone voltage of 50 V, trap collision energy of 4 V, traveling wave ion mobility with a speed of 475 m/s, wave amplitude of 36.5 V, and nitrogen pressure of 2.75 mbar. Low-energy scans applied a transfer collision energy of 4 V, while high-energy scans utilized four distinct collision energy ramps between 15 and 55 V.

ProteinLynx Global Server (PLGS 2.5.1, Waters) was employed to analyse MSE reference data and to identify all detectable peptic peptides. The raw data files were processed, and isotopic distributions assigned using DynamX 3.0 (Waters, USA). The mean values and standard deviations (SD) of these replicates were calculated to assess the significance of changes, with a global significance threshold applied and a T-test (*75*). The peptide level data were also flattened per amino acid (*76, 77*). Custom scripts developed in Python 3.4.0 (Python Software Foundation) were utilized for post-processing analysis, and visualizations were generated using PyMOL (Schrödinger, Inc.).

### Peptide synthesis

Standard Fmoc solid-phase peptide synthesis was performed on a 0.1 mM scale using CEM (Buckingham, UK) Liberty Blue automated peptide synthesis apparatus with inline UV monitoring. Activation was achieved with DIC/Oxyma. Fmoc deprotection was performed with 20% v/v morpholine/DMF. Double couplings were used for β-branched residues and the subsequent amino acid. Peptides were synthesised from C to N terminus as the C-terminal amide on Rink amide resin. For fluorescently labelled peptides TAMRA (0.1 mM, 2 eq.), HATU (0.095 mM, 1.9 eq.) and DIPEA (0.225 mM, 4.5 eq) in DMF (3 mL) were added to DMF washed peptide resin (0.05 mM) with agitation for 3 hours. Resin was washed with 20% piperidine in DMF (5 mL) for 2 x 30 minutes to remove any excess dye. All manipulations were carried out under foil to exclude light. Peptides were cleaved from the solid support by addition of TFA (9.5 mL), TIPS (0.25 mL) and water (0.25 mL) for 3 hours with shaking at rt. The cleavage solution was reduced to approximately 1 mL under a flow of nitrogen. Crude peptide was precipitated upon addition of ice-cold diethyl ether (40 mL) and recovered via centrifugation. The resulting precipitant was dissolved in 1:1 acetonitrile and water (≈ 15 mL) and lyophilised to yield crude peptide as a solid.

### Peptide purification

Peptides were purified by reverse phase HPLC on a Phenomenex (Macclesfield, UK) Luna C18 stationary phase column (150 x 10 mm, 5 μM particle size, 100 A pore size) using a preparative JASCO HPLC system. A linear gradient of 20-80% acetonitrile and water was applied over 30 minutes. Chromatograms were monitored at wavelengths of 220 and 280 nm. The identities of the peptides were confirmed using MALDI-TOF mass spectrometer using a Bruker ultrafleXtreme II instrument in reflector mode. Peptides were spotted on a groundsteel target plate using dihydroxybenzoic acid as the matrix. Peptide purities were determined using a JASCO analytical HPLC system, fitted with a reverse-phase Kinetex® C18 analytical column (Phenomenex, 5 μm particle size, 100 Å pore size, 100 x 4.6 mm). Fractions containing pure peptide were pooled and lyophilised. Peptides were dissolved in buffer (25 mM HEPES pH 7.4 buffer with 150mM NaCl, 5mM BME) and their concentrations determined by UV-Vis on a ThermoScientific (Hemel Hemstead, UK) Nanodrop 2000 spectrophotometer using measurement of UV absorbance at 555 nm (ε_555_(TAMRA) = 85 000 mol^−1^ cm^−1^).

### Circular-dichroism spectroscopy

CD data were collected on a JASCO J-810 or J-815 spectropolarimeter fitted with a Peltier temperature controller (Jasco UK). Peptide samples were dissolved at 100 μM concentration in PBS (8.2 mM sodium phosphate, 1.8 mM potassium phosphate, 137 mM sodium chloride, 2.7 mM potassium chloride at pH 7.4). CD spectra were recorded in 1-mm path length quartz cuvettes at 20 °C. The instruments were set with a scan rate of 100 nm min^−1^, a 1-nm interval, a 1-nm bandwidth and a 1-s response time and scans are an average of 8 scans recorded for the same sample. The spectra were converted from ellipticities (deg) to mean residue ellipticities (MRE; deg.cm^2^.dmol^−1^.res^−1^) by normalising for concentration of peptide bonds and the cell path length using the equation:

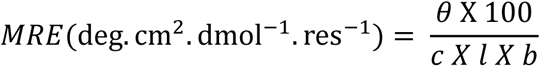

Where the variable *θ* is the measured difference in absorbed circularly polarised light in millidegrees, *c* is the millimolar concentration of the specimen, *l* is the path-length of the cuvette in centimetres and *b* is the number of amide bonds in the polypeptide, for which the N-terminal acetyl bond was included but not the C-terminal amide.

### Sedimentation-equilibrium analytical ultracentrifugation

Analytical ultracentrifugation (AUC) sedimentation-equilibrium experiments were conducted at 20 °C in a Beckman Optima XL-I analytical ultracentrifuge using an An-60 Ti rotor (Beckmann Coulter). Solutions were made up in PBS at 100 μM total peptide concentration. The experiments were run in two-channel centrepiece. The samples were centrifuged at speeds in the range 44–60 krpm and scans at each recorded speed were duplicated. Data were fitted to single, ideal species models using SEDFIT (v15.2b)/SEDPHAT, comprising a minimum of four speeds. 95% confidence limits were obtained via Monte Carlo analysis of the obtained fits.

### Fluorescence polarisation

TAMRA conjugated peptides were diluted to 150 nM and incubated with increasing concentrations of KLC-1/2 TPR protein (typically 0 – 25 μM) in assay buffer (25mM HEPES pH 7.4, 5 mM β-mercaptoethanol and 150mM NaCl). Measurements were performed on a CLARIOstar (BMG Labtech) microplate reader at room temperature. ΔFP values at each concentration were calculated by subtraction of measurements made without TPR protein. Binding constants (*K_d_*) were determined using a one-site specific binding model using GraphPad Prism software. Data were normalized to the calculated B_max_. Where binding was lost or undetectable (e.g. delta Helix1, KinTag-fusion), data were normalised to the B_max_ of the parental protein (e.g. full length KLC1/2 TPR).

### Cell Culture, Immunoprecipitation and Fluorescence Imaging

HeLa cells were grown and maintained in high-glucose Dulbecco’s modified Eagle’s medium (Sigma-Aldrich) supplemented with 10% fetal bovine serum (Sigma-Aldrich) and 1% penicillin/streptomycin (Gibco) at 37°C with 5% CO2. For immunoprecipitation experiments, cells were transfected with plasmids encoding GFP-MAP7 (Addgene plasmid 46076)(*64*) with HA-KLC2 and HA-KIF5C (*47*). Cell lysis and GFP-TRAP immunoprecipitation experiments were performed as described in (*24*). For fluorescence imaging, cells were transfected with HA-KIF5C, KLC2-Halo(*24*), and GFP-MAP7, and labelled with Halo-TMR ligand as previously described (*24*), before fixation and staining with Mouse anti-HA (HA-7) from Sigma-Aldrich, and Alexa 647–conjugated anti-mouse (Thermo Fisher Scientific, and imaging on a confocal microscope.

## Supplementary Figures

**Figure S1.**
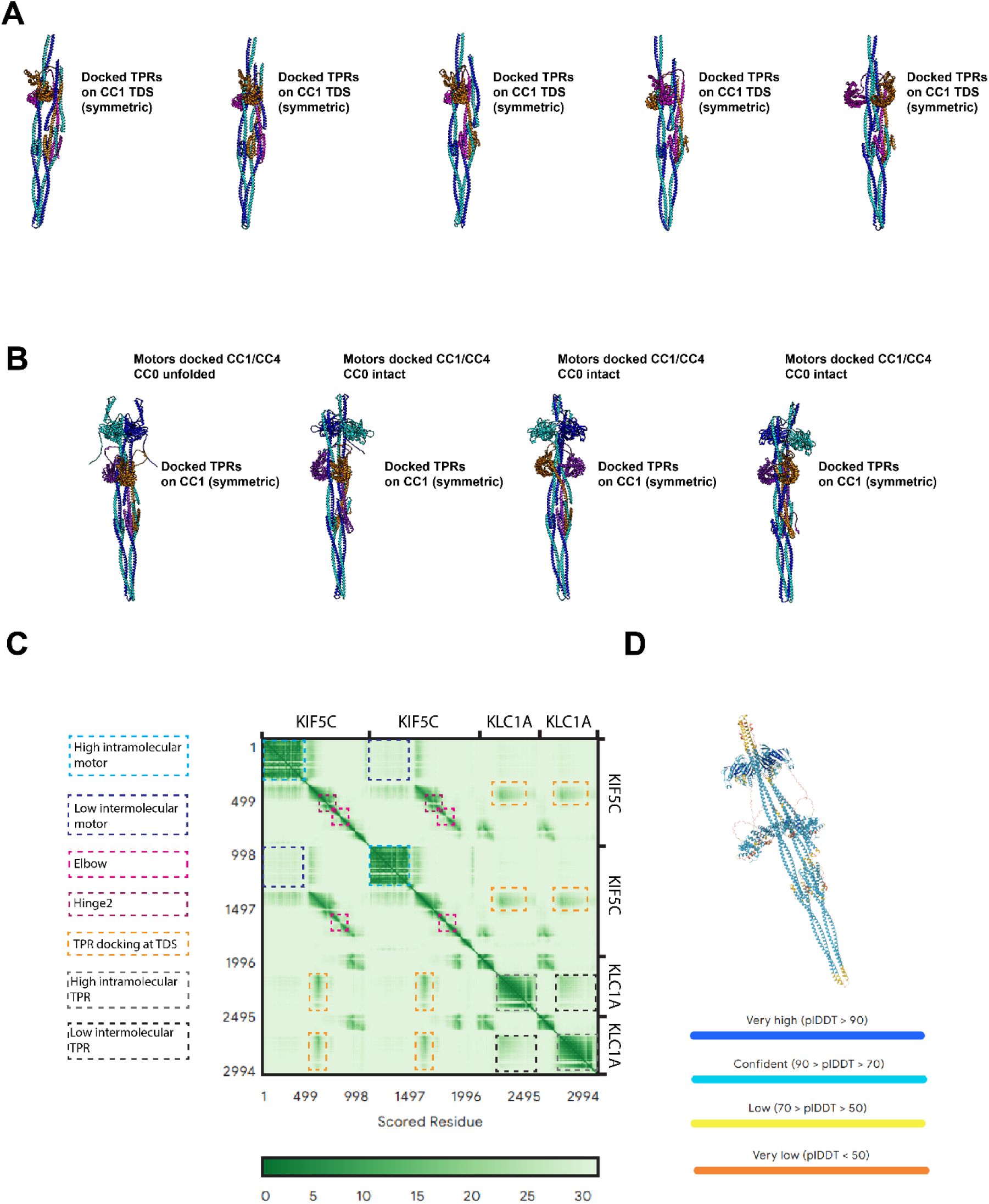
Alphafold3 models of KHC-KLC complexes. (A) Models of kinesin-1 heterotetramer coiled-coil domains (KIF5C CC1-CC4; aa 410-917) with two KLCs (KLC1A CC-TPR; aa 1-480). (B) Models of complete full-length (KIF5C/KLC1A) kinesin-1 heterotetramers, including motor domains and C-terminal KLC sequences. (C) Representative predicted aligned error plot highlighting confidence in intra and inter domain interactions in a full length heterotetrameric model. (D) Representative model showing pLDDT. Plots are typical of other models.

**Figure S2.**
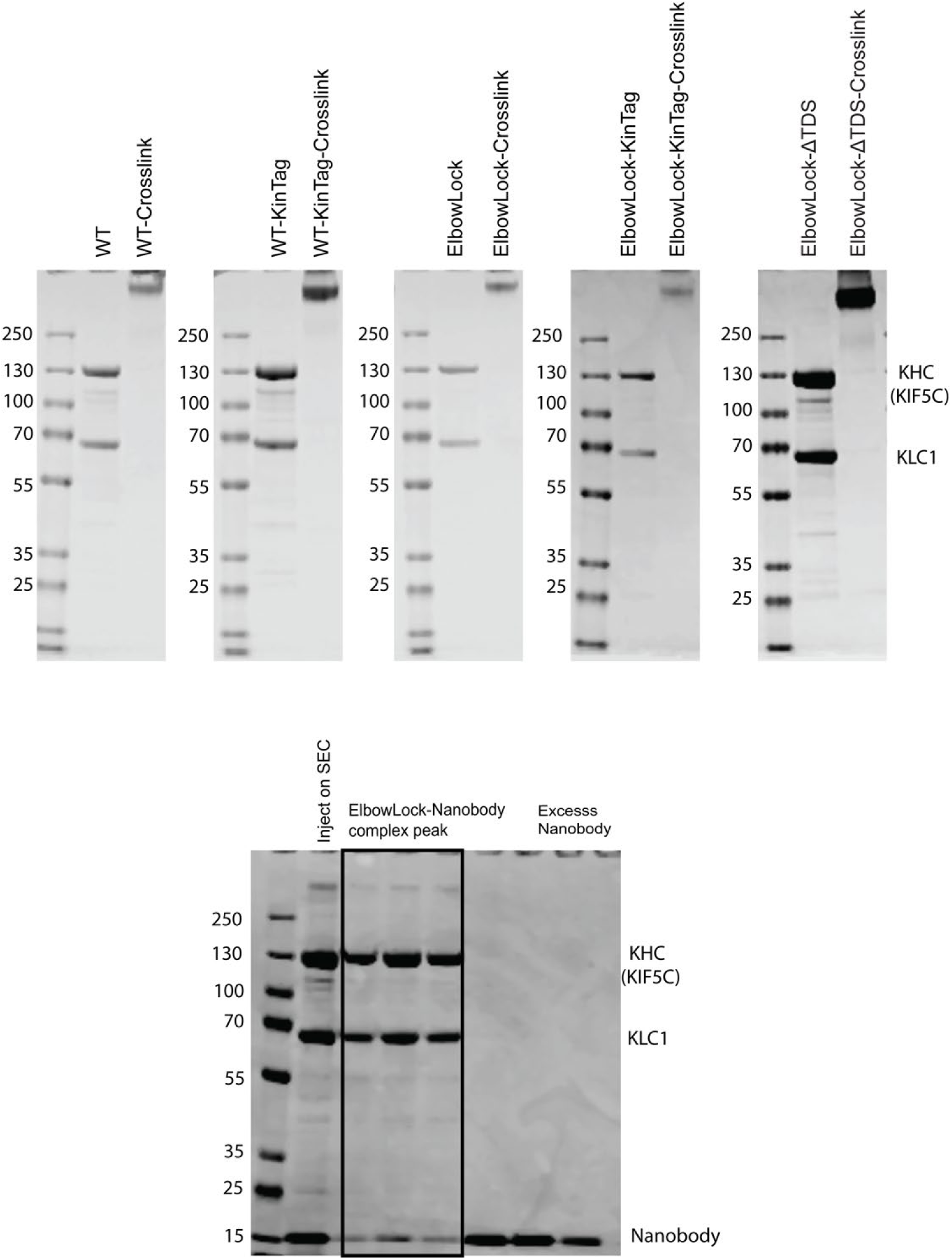
Representative samples of protein complexes used in this study. Coomassie-stained SDS-PAGE gels show samples of proteins after size-exclusion chromatography, with and without BS3 crosslinker (top panel). Coomassie-stained SDS-PAGE gel showing purification of ElbowLock-Nanobody complex from size-exclusion chromatography (bottom panel).

**Figure S3.**
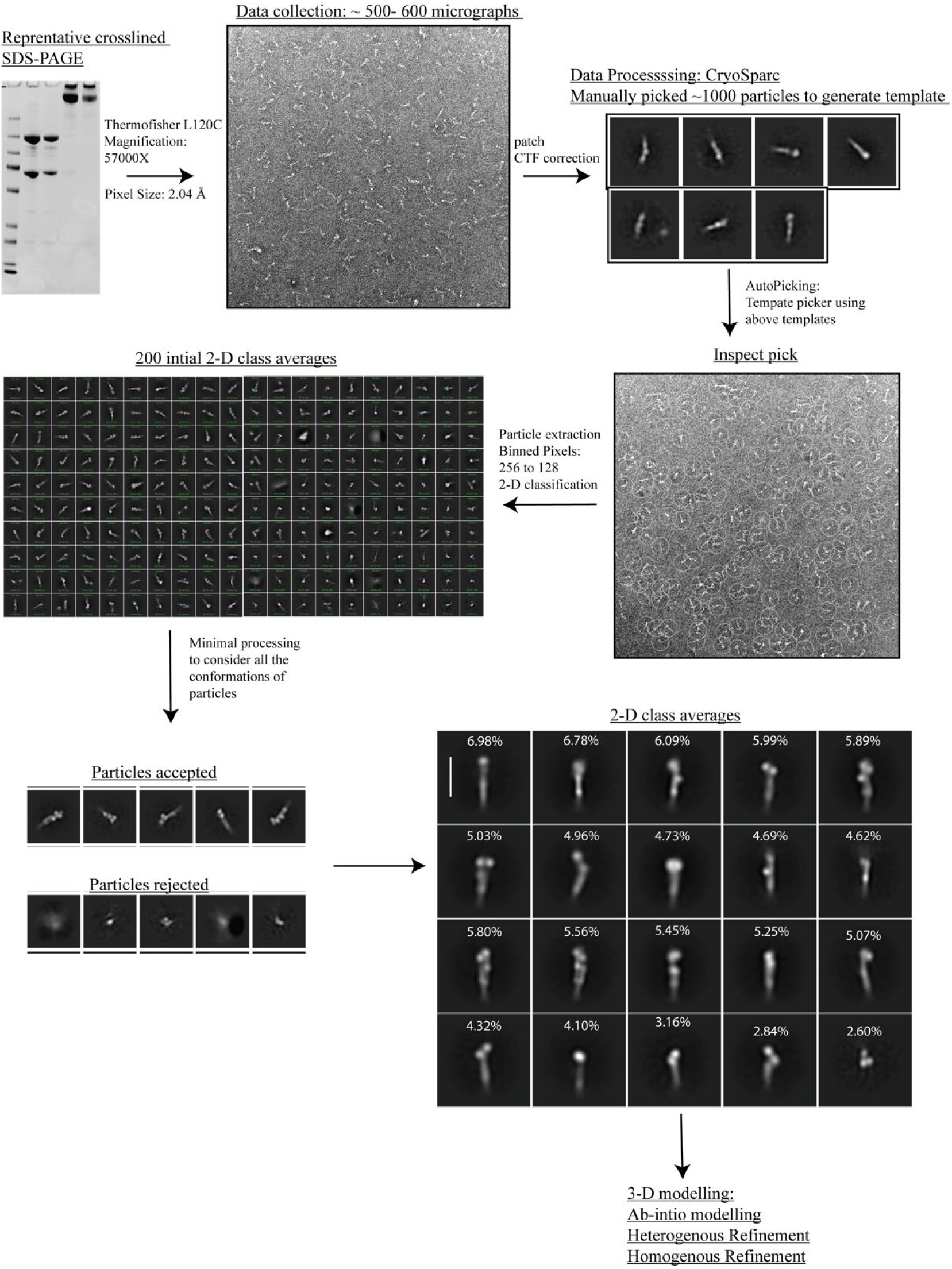
Representative NS-EM processing workflow. Schematic shows typical workflow for 2D classification steps to process NS-EM. Example shown is for ElbowLock complexes.

**Figure S4.**
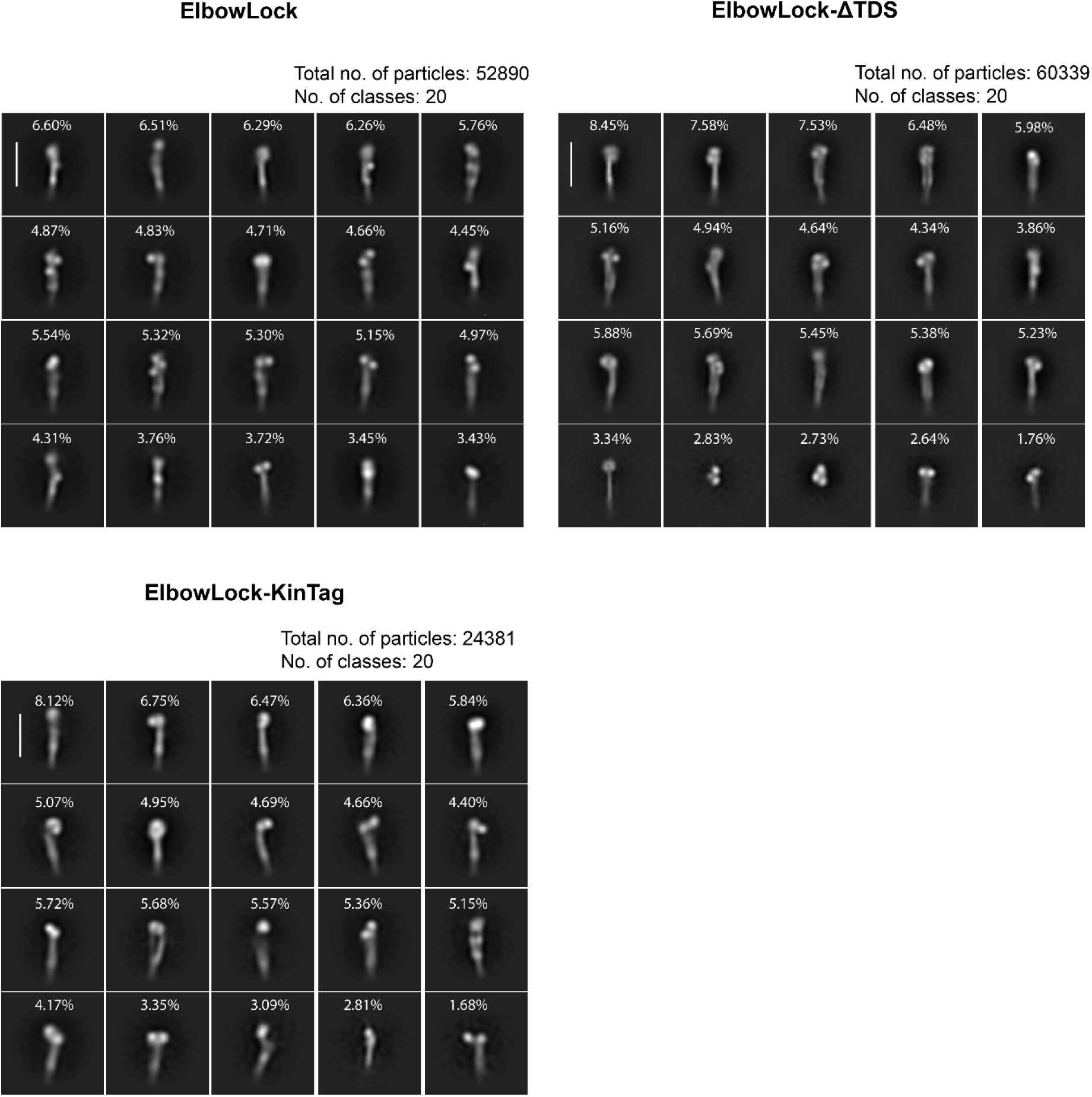
Representative sets of 2D classes for all complexes. The scale bar is 26 nm.

**Figure S5.**
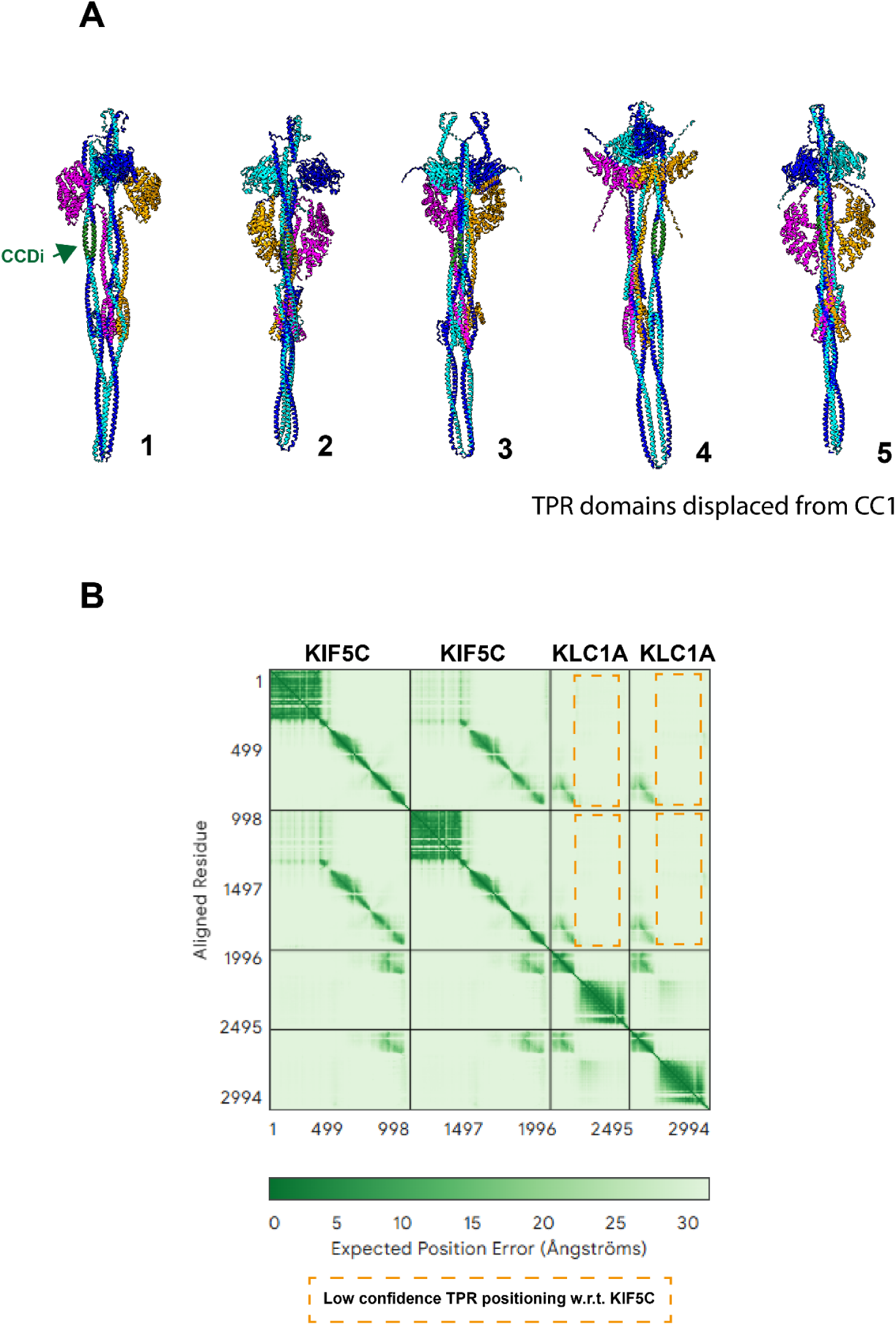
Alphafold3 models of ΔTDS kinesin-1 complexes. (A) 5 models showing ΔTDS complexes (CCDi replaces the TDS sequence showing in green). In all models, the TPR domains are displaced from their original CC1-docked position (compare with Figure 1/S1) and occupy a variety of different positions with very low confidence (B) Representative position error plot (from model 1). Box highlights low confidence in TPR positioning.

**Figure S6.**
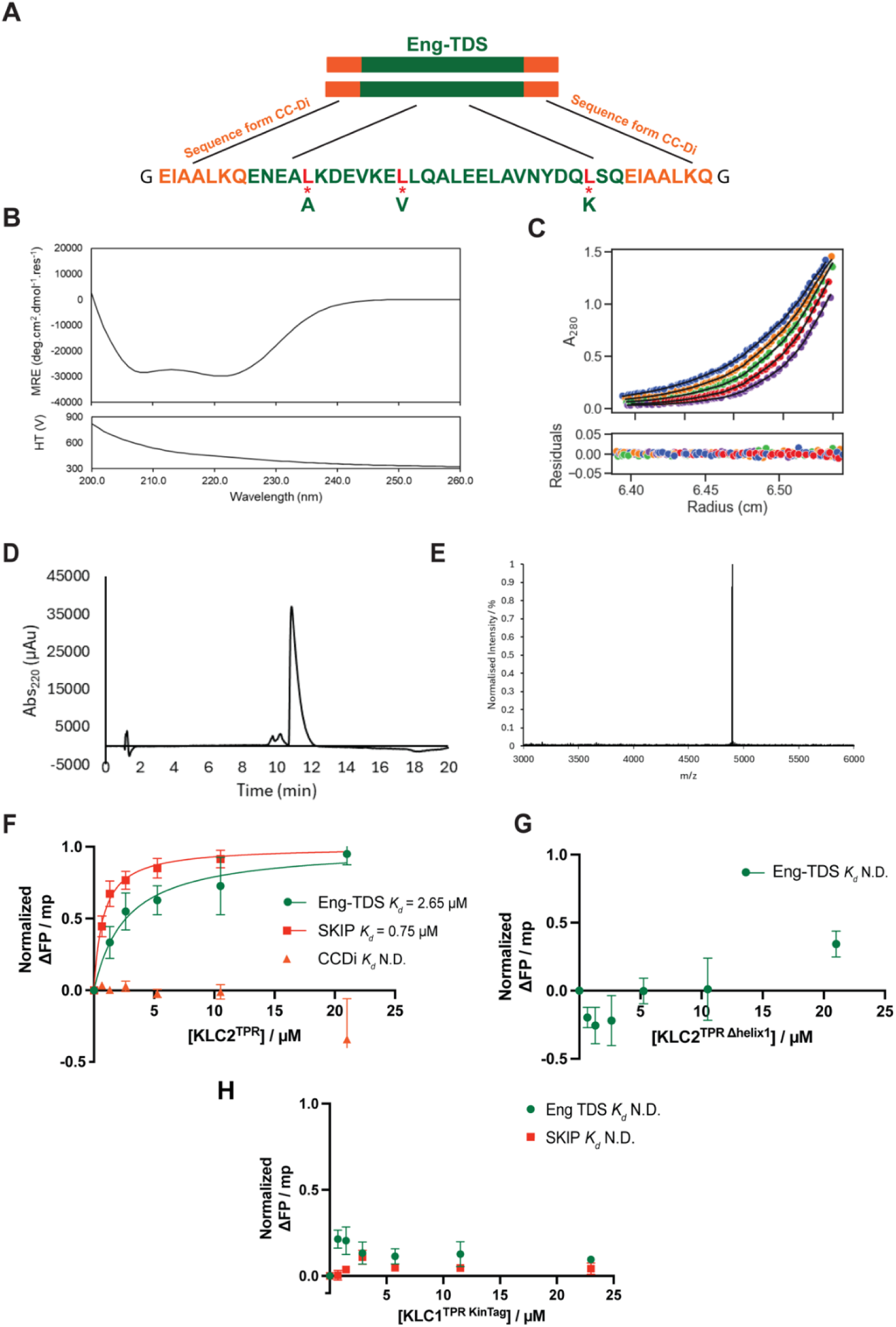
Eng-TDS design, characterisation and interaction with KLC. (A) Schematic showing design of Eng-TDS construct with flanking CCDi sequence and leucine mutations in the hydrophobic core. (B) CD analysis of Eng-TDS indicates alpha helical structure. (C) AUC analysis of Eng-TDS indicates assembly of a dimeric coiled-coil. (D, E) Chromatogram showing the purification of Eng-TDS, accompanied by mass spectrometry analysis confirming a molecular mass of 4895 Da. (F) FP assay showing binding of fluorescently labelled peptides from Eng-TDS and SKIP W-acidic motif to KLC2. (G) FP assay showing no detectable binding of Eng-TDS to KLC2 lacking the first helix of TPR1. (H) Fluorescence polarisation binding assays show no detectable binding of fluorescently labelled TDS and SKIP (W-acidic motif) peptides to isolated KLC1-TPR-KinTag fusion proteins.

**Figure S7.**
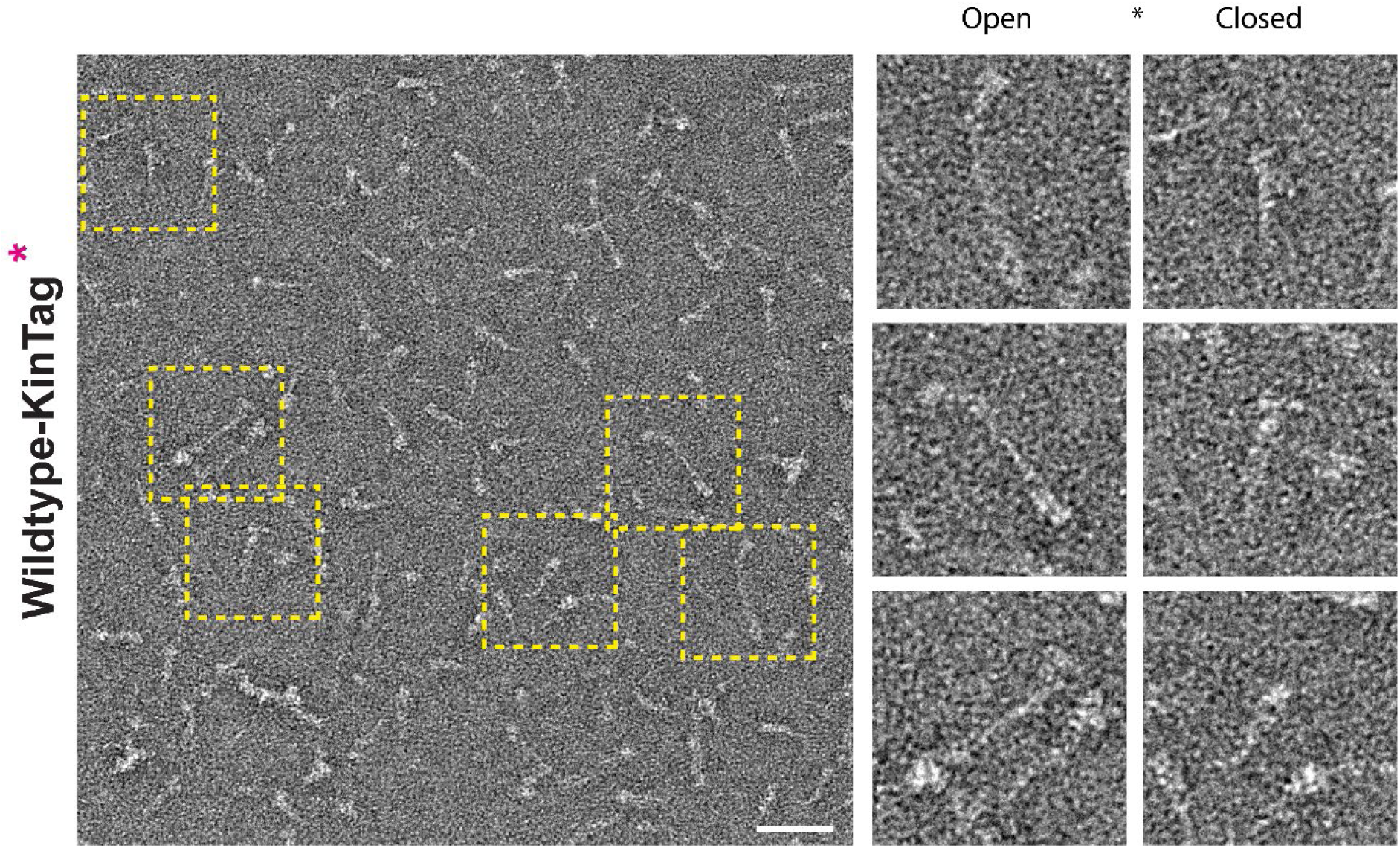
Representative NS-EM electron micrograph showing wild type-KinTag particles with selected open and close particles expanded on right. The scale bar is 50 nm.

**Figure S8.**
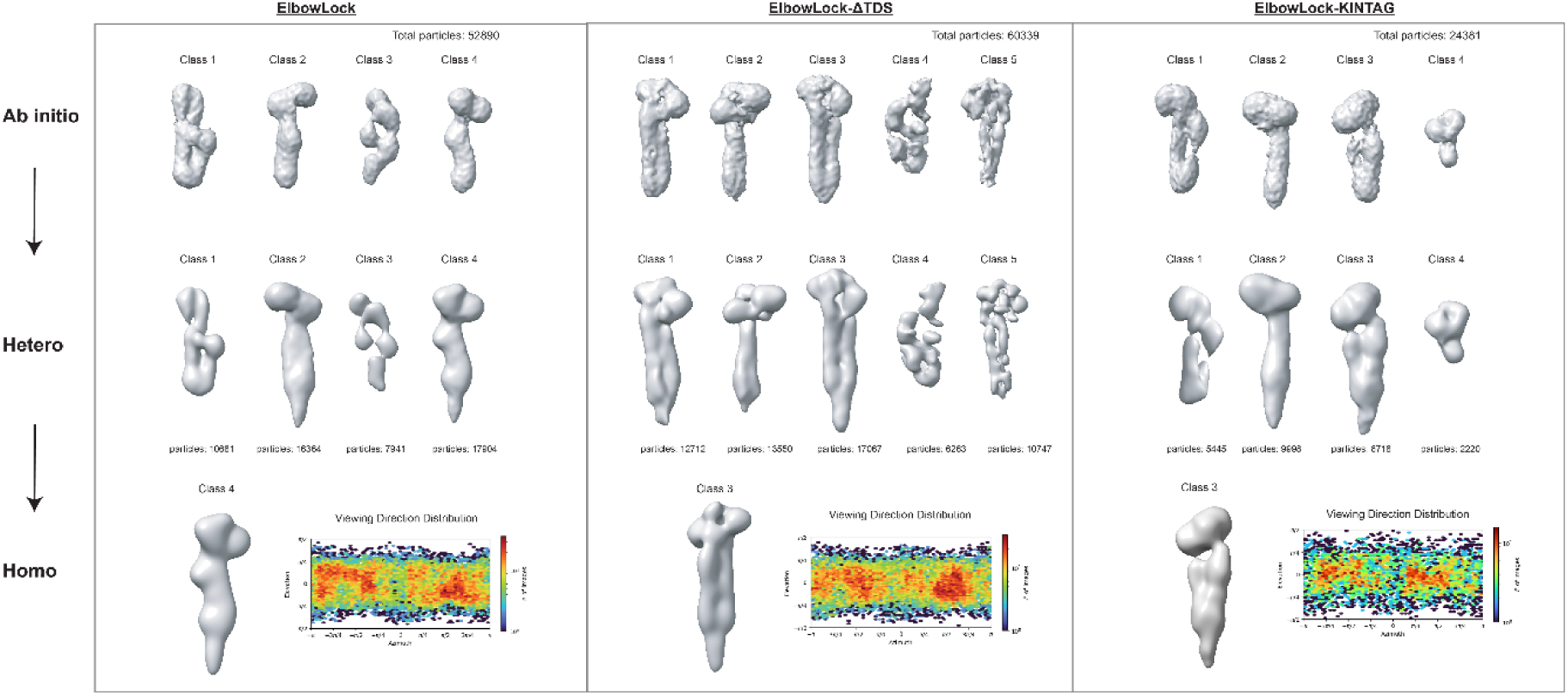
EM processing workflow for 3D reconstructions

**Figure S9.**
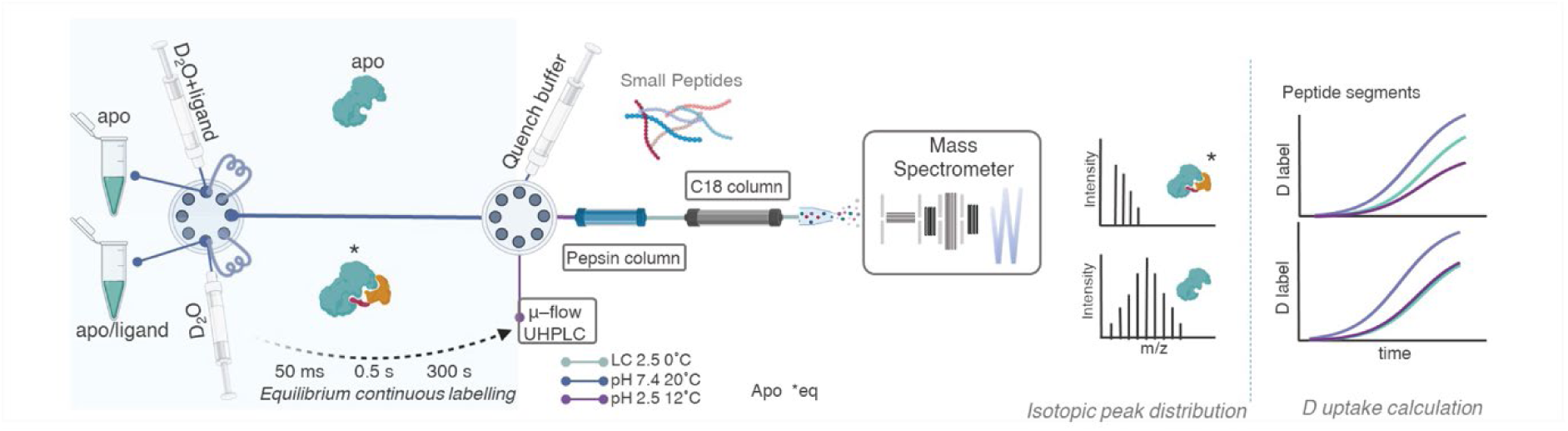
Schematic representation of the HDX experiment.

**Figure S10.**
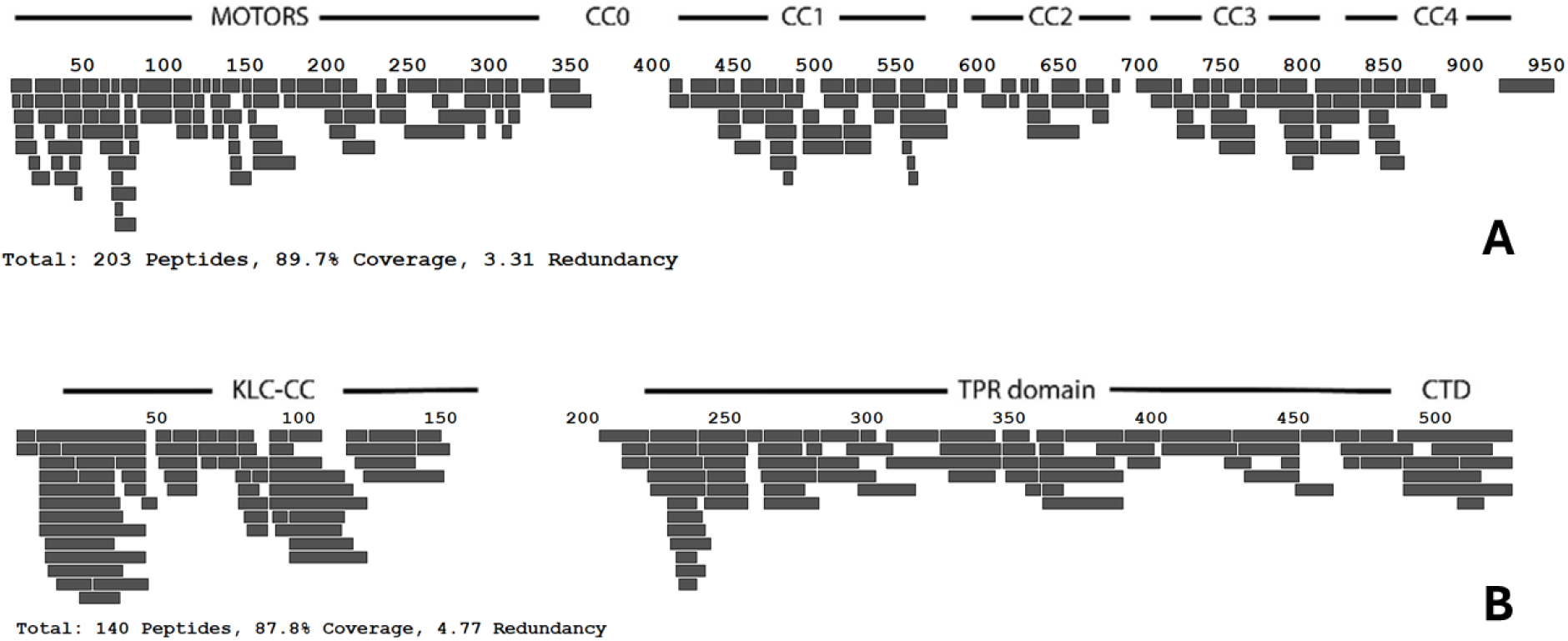
Coverage map of mass spectral assignment of peptides in ‘bottom-up’ HDX-MS experiments for kinesin-1 HC (A) and LC (B).

**Figure S11.**
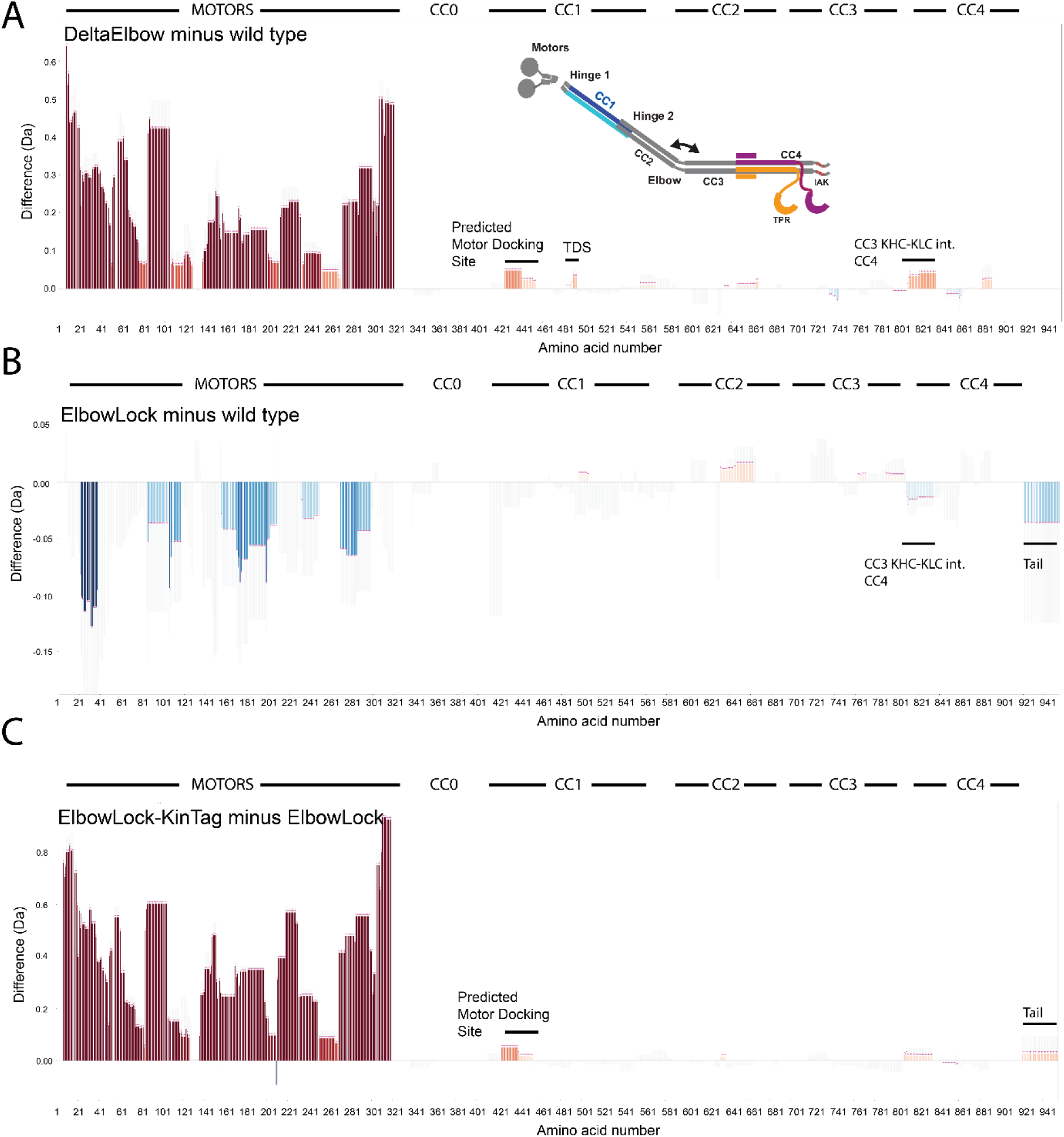
Difference plots showing total HDX across timepoints comparing heavy chain of DeltaElbow and Elbowlock to wild type and ElbowLock-KinTag to ElbowLock. The difference in the total observed deuterium uptake at 0.3, 0.5, 1, 3, 10, 30, and 300 seconds was summed and compared between DeltaElbow and wild type (A) and ElbowLock and wild type (B), and ElbowLock-KinTag to ElbowLock (C). In each case, the data for the later were subtracted from the former. Therefore, relative protection in a region of the former results in a more negative value (blue bars extending downward), while deprotection, such as from an exposed domain interface, results in a more positive value (red bars extending upward). Each vertical bar represents a distinct peptide, and the horizontal axis corresponds to amino acid number from N to C. Significant differences in deuterium labelling per peptide and time point, derived from a T-test, are marked with pink asterisks.

**Figure S12.**
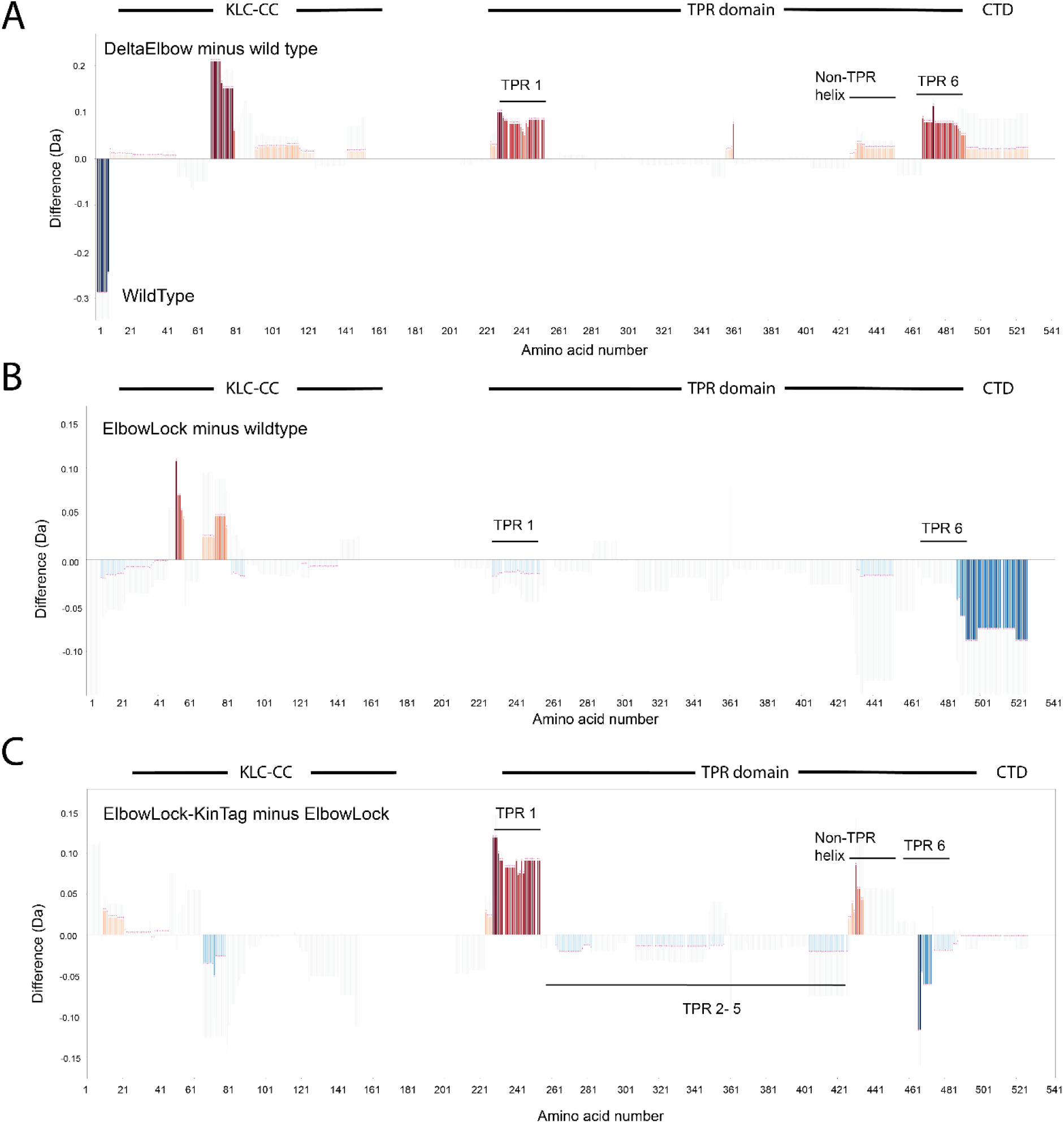
Difference plots showing total HDX across all timepoints comparing light chain of DeltaElbow and Elbowlock to wild type and ElbowLock-KinTag to ElbowLock. The difference in the total observed deuterium uptake at 0.3, 0.5, 1, 3, 10, 30, and 300 seconds was summed and compared between DeltaElbow and wild type (A) and ElbowLock and wildtype (B), and ElbowLock-KinTag to ElbowLock (C). In each case, the data for the later were subtracted from the former. Therefore, relative protection in a region of the former results in a more negative value (blue bars extending downward), while deprotection, such as from an exposed domain interface, results in a more positive value (red bars extending upward). Each vertical bar represents a distinct peptide, and the horizontal axis corresponds to amino acid number from N to C. Significant differences in deuterium labelling per peptide and time point, derived from a T-test, are marked with pink asterisks.

**Table S1.**
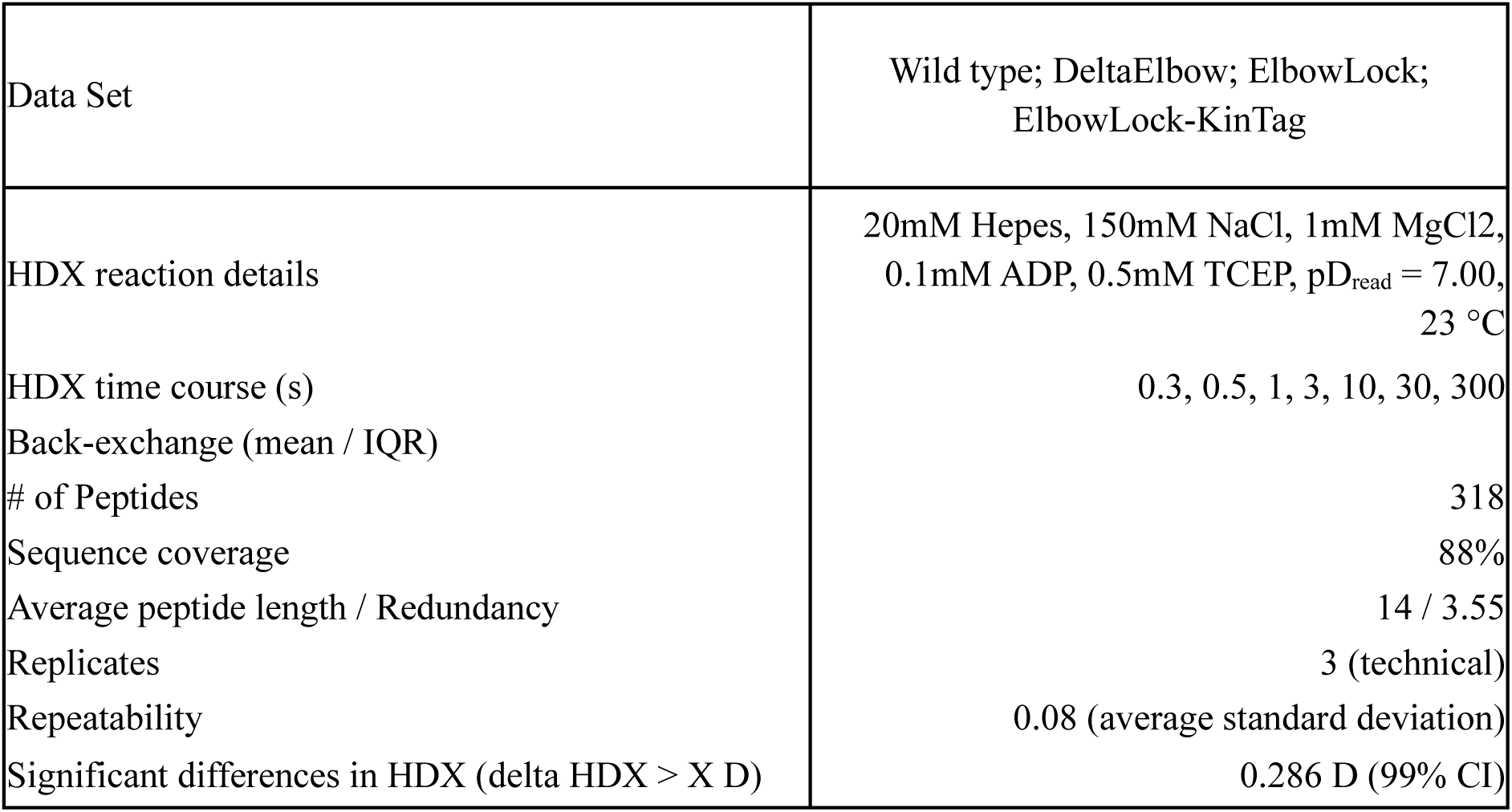
HDX summary table.

| Data Set | Wild type; DeltaElbow; ElbowLock;<br>ElbowLock-KinTag |
| --- | --- |
| HDX reaction details | 20mM Hepes, 150mM NaCl, 1mM MgCl<sub>2</sub>,<br>0.1mM ADP, 0.5mM TCEP, pD<sub>read</sub> = 7.00,<br>23 °C |
| HDX time course (s) | 0.3, 0.5, 1, 3, 10, 30, 300 |
| Back-exchange (mean / IQR) |  |
| # of Peptides | 318 |
| Sequence coverage | 88% |
| Average peptide length / Redundancy | 14 / 3.55 |
| Replicates | 3 (technical) |
| Repeatability | 0.08 (average standard deviation) |
| Significant differences in HDX (delta HDX > X D) | 0.286 D (99% CI) |

